# Cohort-based smoothing methods for age-specific contact rates

**DOI:** 10.1101/290551

**Authors:** Yannick Vandendijck, Oswaldo Gressani, Christel Faes, Carlo G. Camarda, Niel Hens

**Affiliations:** Interuniversity Institute for Biostatistics and statistical Bioinformatics (IBioStat), Hasselt University, Belgium; French Institute for Demographic Studies (INED), Aubervilliers, France; Centre for Health Economics Research and Modelling Infectious Diseases, Vaxinfectio, University of Antwerp, Belgium

**Keywords:** Penalized iterative reweighted least squares, Penalized likelihood, Constrained smoothing, Social contact matrix

## Abstract

The use of social contact rates is widespread in infectious disease modeling since it has been shown that they are key driving forces of important epidemiological parameters. Quantification of contact patterns is crucial to parametrize dynamic transmission models and to provide insights on the (basic) reproduction number. Information on social interactions, can (for instance) be obtained from population-based contact surveys, such as the European Commission project POLYMOD. Estimation of age-specific contact rates from these studies is often done using a piecewise constant approach or bivariate smoothing techniques. For the latter, typically, smoothing is done in the dimensions of the respondent’s and contact’s age. We propose a new flexible strategy based on a smoothing constrained approach - taking into account the reciprocal nature of contacts - where the contact rates are assumed smooth from a cohort perspective as well as from the age distribution of contacts. This is achieved by smoothing over the diagonal components (including all subdiagonals) of the social contact matrix. This approach is supported by the fact that people age with time and thus motivates smoothly varying contact rates from a cohort angle. Two approaches that allow for smoothing of social contact data over cohorts are proposed namely, (1) reordering of the diagonal components of the social contact matrix; and (2) reordering of the penalty matrix associated with the diagonal components. Parameter estimation is done in the likelihood framework by using constrained penalized iterative reweighted least squares (C-PIRLS), under Poisson and negative Binomial distributional assumptions for the observed contacts. A simulation study underlines the benefits of cohort-based smoothing based on two scalar measures of performance. Finally, the proposed methods are illustrated on the Belgian POLYMOD data of 2006. Code to reproduce the results of the article can be downloaded on this Github repository https://github.com/oswaldogressani/Cohort_smoothing.

## 1 Introduction

Understanding the spread of infectious diseases in an epidemic context is a challenging task for mathematical modelers. It is especially made difficult by the complexities and intricacies of demography dynamics and rich social contact networks. Social contact mixing patterns play a key role in assessing disease transmission and are known to be crucial determinants of important epidemiological parameters such as the basic reproduction number and the force of infection (see e.g., Vynnycky and White, 2010; Hens et al., 2009). One approach to account for mixing patterns is by the use of the so-called “Who Acquires Infection From Whom” (WAIFW) matrix and the use of serological data to estimate the WAIFW parameters (Anderson and May, 1991; Greenhalgh and Dietz, 1994; Farrington et al., 2001; Van Effelterre et al., 2009). Another approach proposed by Farrington and Whitaker (2005) is to model contact rates as a continuous surface and estimate parameters from serologic survey data. The main limitations of both approaches is that they rely on structural assumptions on the WAIFW matrix and on an arbitrary choice of the parametric model used for the continuous contact surface. Alternatively, over the last two decades or so, several studies have reported on ways of collecting data on social mixing behaviour relevant to the spread of close contact infections directly from individuals through self-reported number of contacts (Wallinga et al., 2006; Beutels et al., 2006; Edmunds et al., 1997, 2006; Mikolajczyk et al., 2007). The POLYMOD initiative can arguably be counted among the most important studies in infectious disease epidemiology in Europe, providing a large and representative popoulation based survey on social contacts (Mossong et al., 2008). The estimation of smooth age-specific contact rates from the POLYMOD project data is typically performed by applying a negative Binomial model on the aggregated number of contacts. To ensure enough flexibility, a bivariate frequentist smoothing method is implemented by using a tensor product spline as a function of the respondent’s and contact’s ages as a smooth interaction term (Mossong et al., 2008; Hens et al., 2009; Goeyvaerts et al., 2010). From a Bayesian perspective, van de Kassteele et al., 2017 estimate social contact rates by means of a Gaussian Markov Random Field (GMRF) and use Integrated Nested Laplace Approximations (INLA) Rue et al. (2009) as the main tool for inference.

We propose a new smoothing constrained approach, where contact rates are assumed to be smooth both from a cohort perspective and from the age distribution of contacts. This means that smoothing in the direction of the age of contacts will remain. However, smoothing over the dimension of the age of respondents will be replaced by smoothing contact rates from a cohort perspective by focusing on the diagonal compontents (including all subdiagonals) of the social contact matrix. Under the likelihood framework and assuming Poisson or negative Binomial models for the aggregated number of contacts, diagonal smoothing of contact matrices is achieved through two alternative approaches: (1) reordering of the diagonal components yielding a rectangular grid; and (2) reordering of the penalty matrix to translate a penalization scheme over the diagonal components. The first approach builds further upon work published by two of the co-authors in a proceedings paper (*reference not provided in reviewing process*).

The article is organized as follows. Section 2 aims at presenting three competing approaches to smoothly estimate social contact rates. Section 3 investigates the statistical performance of the proposed approaches through a numerical study and Section 4 illustrates the methodology on the Belgian POLYMOD data. Finally, Section 5 concludes with a discussion and prospects for future research.

## 2 Smoothing social contact data

In this section, we present three competing smoothing constrained approaches (SCAs) to infer social contact rates. First, we describe the classic approach where smoothing is performed in the dimensions of the respondent’s and contact’s ages, thus ignoring the cohort effect. The latter baseline model will be referred to 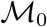. Second, we present the new competing models, namely the SCA where contact rates are assumed smooth from a cohort perspective. Two approaches are investigated both in terms of performance and computational speed, namely model 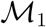, where a reordering of the diagonal components is considered to reproduce a rectangular contact matrix; and model 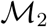, where a reordering of the components of the penalty matrix yields a penalization scheme targeting the diagonal components of the social contact matrix.

### 2.1 Absence of smoothing over cohorts

Let **Y** = (*y_ij_*) be a square (*m* × *m*) matrix where the *ij*th entry is the total number of contacts made by the respondents of age *i* – 1 with individuals of age *j* – 1, with indices *i* = 1,…, *m* and *j* = 1,…, *m*. This information can be extracted from the selfreported contact diaries of the participants and in our specific case *m* = 77 for the Belgian POLYMOD data. Let **y** be the *m*^2^ × 1 vector obtained by arranging the matrix **Y** by row order into a vector. Furthermore, let the *m* × 1 vector **r** = (*r_i_*) contain the total number of respondents of age *i* – 1. Define the *m* × *m* matrix **E** = **r1**_*m*_, where **1**_*m*_ is a 1 × *m* vector of ones, and define **e** as the *m*^2^ × 1 vector obtained by arranging the matrix **E** by row order into a vector. Let the *m* × 1 vector **p** = (*p_i_*) denote the population size of individuals of age *i* – 1 and define the *m* × *m* matrix **P** = **p1**_*m*_. The supplementary materials provide examples of how to construct these vectors and matrices for the specific case *m* = 4. The expected number of contacts made by participants of age *i* – 1 with contacts of age *j* – 1 is denoted by *E*(*y_ij_*) = *μ_ij_* = *r_i_γ_ij_*, where *γ_ij_* is the actual contact rate of individuals of age *i* – 1 with contacts of age *j* – 1. In other words, *γ_ij_* is the average number of contacts an individual of age *i* – 1 makes with an individual of age *j* – 1. Define the so-called social contact matrix **Γ** as the *m* × *m* matrix with elements *γ_ij_* (see Figure 1 left panel) and let ***γ*** be the *m*^2^ × 1 vector obtained by arranging the matrix **Γ** by row order into a vector.

**Figure 1:**
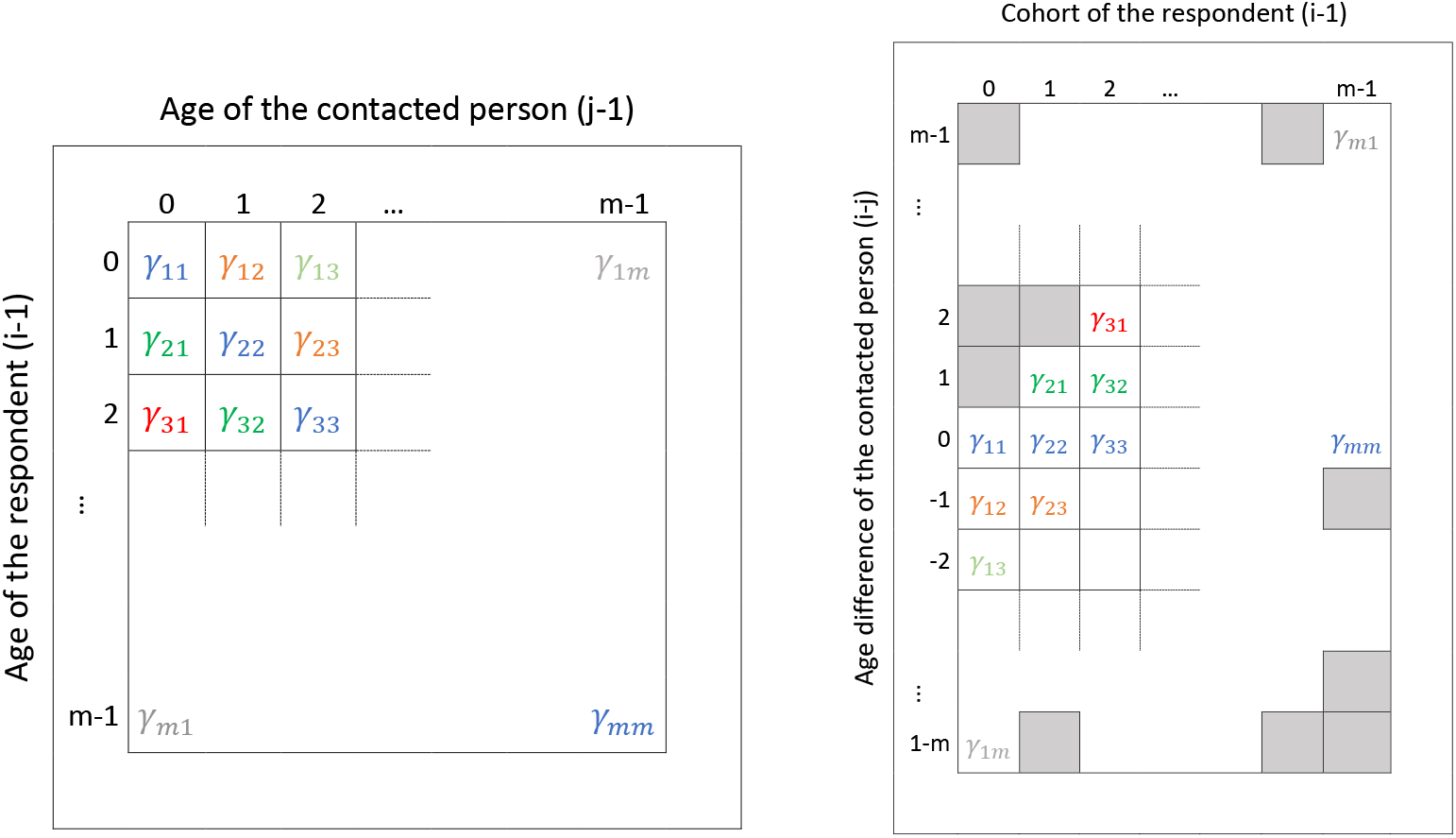
Schematic representation of the original data structure of **Γ** over ages of respondents and ages of contacts (left panel) and the restructured matrix 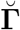 over cohorts of the respondents and age differences of the contacted persons (right panel). Cells with nuisance parameters in 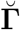 are depicted with gray squares.

The expected number of contacts can also be written as E(**y**) = ***μ*** = **e**⊙***γ***, where ⊙ denotes component-wise multiplication (also known as the Hadamard product). The interest lies in estimating the unknown contact parameters *γ_ij_* from data **y** in a smooth way, such that the important signal in the mixing patterns is captured. For this purpose, we assume that the observed contacts (*y_ij_*) are realizations from a Poisson distribution, i.e. **y** ~ Poiss(***μ***). For modeling purposes, a log-link function is specified, namely log(***γ***) = ***η***, so that log(***μ***) = log(**e**) + log(***γ***) = log(**e**) + ***η***.

Let **H** be the *m* × *m* matrix with *ij*th element *η_ij_*. Interest is in the estimation of the *m*^2^ unknown parameters ***η***. It can be readily seen that the maximum likelihood estimates are given by 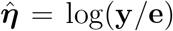, and thus 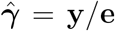, in case the parameters can be estimated freely. However, these estimates do not yield a smooth contact rate surface and hence are only of interest for exploratory purposes. We prefer to work with a modeling approach that yields social contact rates that are smooth and reciprocal. Reciprocity of contacts in this context means that the total number of contacts on the population level from age *i* to age *j* must equal the total number of contacts from age *j* to age *i*. The latter reciprocal nature can be expressed mathematically as *γ_ij_p_i_* = *γ_ji_p_j_* for all *i* = 1,…, *m* and *j* = 1,…, *m* and can be written as the difference log(*γ_ij_*) – log(*γ_ji_*) = log(*p_j_*) – log(*p_i_*) and thus:

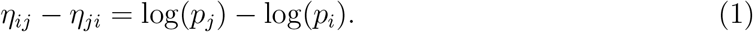

In matrix form:

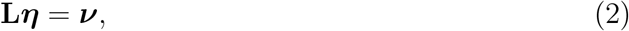

where **L** is a 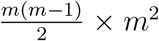 allocation matrix with entries +1 and −1 to suit the left-hand side of (1) and vector ***ν*** is given by:

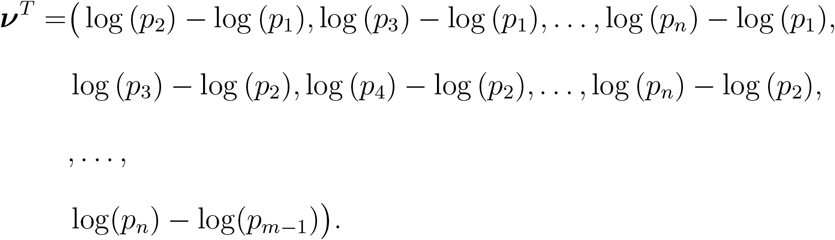

Estimation of the smoothed parameters ***η*** that satisfy the reciprocal constraints is performed through constrained penalized iterative reweighted least squares (C-PIRLS) (Nelder and Wedderburn, 1972; McCullagh and Nelder, 1989; Eilers and Marx, 1996; Wood, 2006). Given current estimates 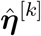 and 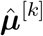 at iteration *k*, parameter estimates 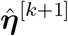 at iteration *k* + 1 are obtained by solving the set of linear equations:

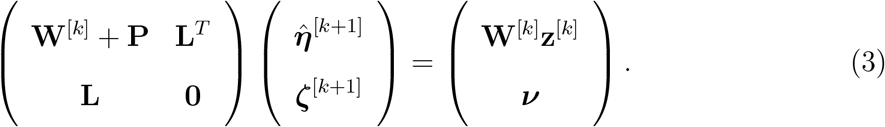

The parameter estimates 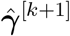 are obtained by exponentiation 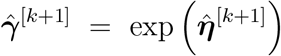. In (3), ***ζ***^[*k*+1]^ is a 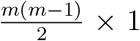 vector of Lagrange multipliers, **W**^[*k*]^ is a *m*^2^ × *m*^2^ diagonal matrix with entries 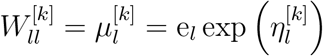 and **z**^[*k*]^ is a *m*^2^ × 1 vector of the so-called *pseudodata* given by:

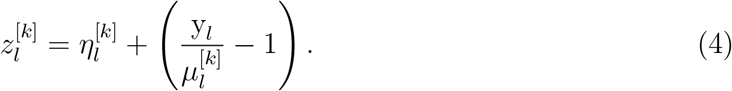

To enforce smoothness over two dimensions, the penalty term **P** in (3) is a *m*^2^ × *m*^2^ matrix given by (see Marx and Eilers, 2005):

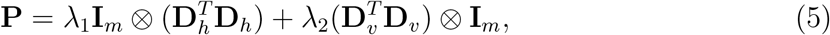

where ⊗ denotes the Kronecker product and λ_1_ and λ_2_ are smoothing parameters for, respectively, the horizontal and vertical dimension in Figure 1 (left panel). The matrices **D**_*h*_ and **D**_*v*_ are second order difference matrices and **I** is the identity matrix. The above iterative process is repeated until convergence, namely until max 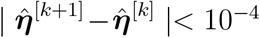. The optimal smoothing parameters are chosen based on minimization of the Akaike Information Criterion (Akaike, 1973) via a grid search:

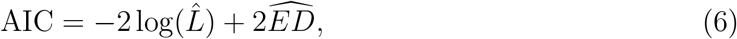

where 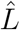 is the maximized value of the likelihood function and the effective degrees of freedom, 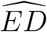, is the trace of the hat matrix given by (see Wood, 2006):

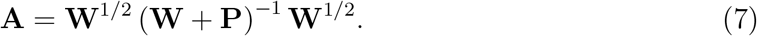

### 2.2 Cohort smoothing by reordering the contact matrix (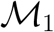 model)

In the previous section, contact rate parameters are smoothed in the dimensions of the respondent’s and contact’s ages. We now describe a new strategy where contact rates are smoothed over the diagonal components and thus over cohorts. In addition, we also smooth over the dimension of the contact’s age since the distribution of the age of (grand)parents can in general be assumed smooth (e.g., children will meet their parents and grandparents who are, for example, ± 30 and ±60 years older). We describe how this can be achieved by restructuring the data and contact matrix over the cohorts and the contacts’ ages.

The contact matrix **Γ** is restructured in such a way that each diagonal (the main diagonal and all sub-diagonals) is present as a row in the restructured matrix. The restructured matrix is denoted 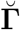. Figure 1 (right panel) gives a graphical representation of this restructured matrix. The matrix 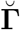 has dimension (2*m* – 1) × *m* and is constructed by entering row *i* of **Γ** in column *i* of 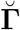 at positions *m* – *i* + 1 to 2*m* – *i*. In that manner, all subsequent diagonal elements are present in the same row. By construction, matrix 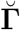 contains nuisance contact rate parameters that are not directly of interest. Restructured matrices 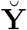 and **Ĕ**, constructed from **Y** and **E**, are created similarly as 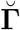. Missing cell entries are present for 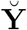 and **Ĕ** at the same cells where the nuisance parameters are present for 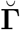. To handle these missing entries, we impute arbitrary values (say, 9999) in 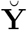 and **Ĕ** and construct a (2*m* – 1) × *m* weight matrix 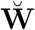, where the *ij*th entry of 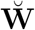 equals zero if the *ij*th entry in 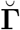 is a nuisance parameter and equals one otherwise. This weight matrix avoids that the imputed values for the missing entries influence parameter estimation.

Let 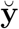, **ĕ**, 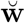 and 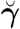 be the (2*m*^2^ – *m*) × 1 vectors obtained by arranging the matrices 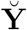, **Ĕ**, 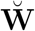 and 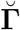 by column order into a vector. Again, we assume that 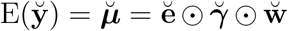 and that the observations come from a Poisson distribution, namely 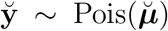 and 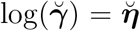. The objective is now to estimate the 2*m*^2^ – *m* unknown parameters 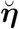. However, only the *m*^2^ parameters of 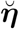 corresponding to the non-nuisance parameter entries in 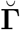 are of interest. The reciprocity assumption of the contacts, namely 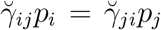, can again be written in matrix form as:

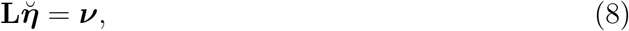

where **L** is an 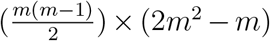 allocation matrix to accommodate the reciprocity constraints. Estimation of the smoothed parameters 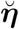 is again performed through C-PIRLS. Updated parameter estimates are now obtained by solving the set of linear equations:

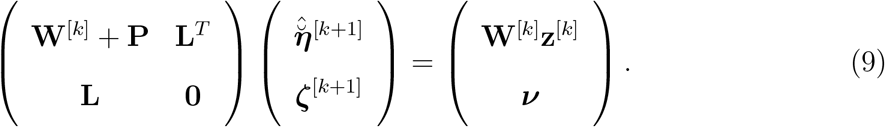

In (9), ***ζ***^[*k*+1]^ are again Lagrange multipliers, **W**^[*k*]^ is an (2*m*^2^ – *m*) × (2*m*^2^ – *m*) diagonal matrix with entries 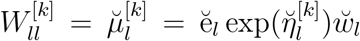 and **z**^[*k*]^ is an (2*m*^2^ – *m*) × 1 vector of pseudovalues given by:

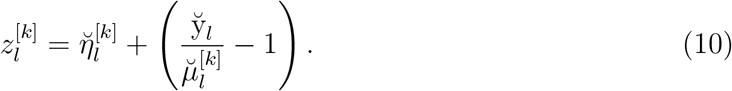

Here, the penalty term **P** in (9) is a (2*m*^2^ – *m*) × (2*m*^2^ – *m*) matrix given by:

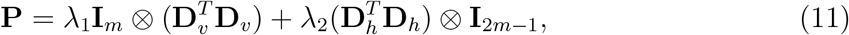

where λ_1_ and λ_2_ are smoothing parameters for, respectively, the vertical and horizontal dimension in Figure 1 right panel, i.e. age and cohort of the original data structure. Optimal smoothing parameters are again computed via grid search using the AIC.

### 2.3 Cohort smoothing by reordering the penalty matrix (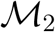 model)

An altenative approach to smooth over cohorts is to work from the perspective of the penalty matrix without rearranging the original social contact matrix. The methodology is very similar as the one described in Section 2.1. Matrices **Y**, **E**, **P**, **Γ** and vectors **y**, **e**, **μ**, **γ**, **η** are defined similarly as in Section 2.1. Again, a Poisson distribution is assumed for the observed contact rates and the reciprocity constraint is written in matrix form as **L*η*** = ***ν***. C-PIRLS is used once more to solve the set of linear equations in (3) for parameter estimation. The penalty matrix **P**, constructed differently as the penalty term in (5), is now a *m*^2^ × *m*^2^ matrix given by:

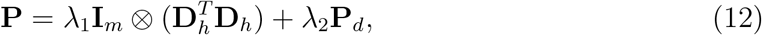

where λ_1_ and λ_2_ are smoothing parameters for, respectively, the horizontal and the diagonal dimension in Figure 1 (left panel), with optimal values chosen by the AIC. The *m*^2^ × *m*^2^ matrix **P**_*d*_ is responsible for the penalization of the parameters of the cohorts (all diagonals and subdiagonals). For example, in the specific case where **Γ** is a 4 × 4 matrix (i.e. ***γ*** = {*γ*_11_, *γ*_12_, *γ*_13_, *γ*_14_, *γ*_21_,…, *γ*_44_}), the penalty matrix **P**_*d*_ is a 16 × 16 matrix (see appendix A).

A major advantage of using the penalty matrix **P**_*d*_ to achieve cohort smoothing is the absence of nuisance parameters in the matrix **Γ** (cf. the approach in the previous section using 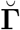). This entails a non-negligible computational gain, since only *m*^2^ parameters in **Γ** need to be estimated, whereas the 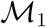 approach requires estimation of 2*m*^2^ – *m* parameters in 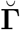 (thus including *m*^2^ – *m* nuisance parameters). However, the modified penalty matrix **P**_*d*_ is less trivial to construct. Whereas the penalty in (11) is easily obtained using standard matrix multiplication, the construction of **P**_*d*_ requires an algorithm (see supplementary materials). This is not a major disadvantage as the construction of **P**_*d*_ is performed only once and outside the C-PIRLS algorithm without requiring a too high computational budget.

### 2.4 Kink on the main diagonal of the social contact matrix

The use of smoothing approaches for estimating social contact rates can lead to estimates that are oversmoothed for individuals of the same age, meaning that the estimated contact rate is smaller than the true one in the population. For example, students make an above average number of contacts with individuals of their own age (e.g., in school, sport clubs, etc.). Smoothing approaches thus, potentially, lead to an underestimation of the social contact rates on the main diagonal of the contact matrix, especially for children and young adults. To take this into account, we introduce the use of a so-called *kink* on the main diagonal of the social contact matrix for 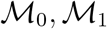 and 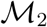, that can force a sudden increase (or decrease) of the estimated social contact rates for children and young adults of the same age.

The kink is introduced through a small adjustment in the penalty matrices in (11) and (12). More specifically, in the dimension of the contact’s age, the social contact rates that belong to the main diagonal, i.e. *η_ii_* and *γ_ii_*, are not penalized. In (11) this is achieved by changing the (2*m* – 3) × (2*m* – 1) matrix **D**_*v*_ as follows:

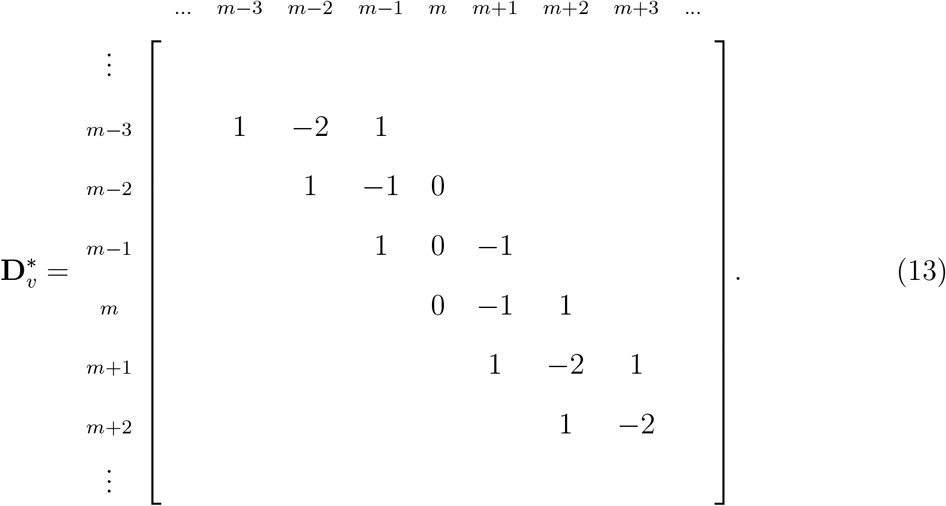

From the above matrix 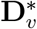, it is clear that the social contact rates that belong to the main diagonal, namely, *η_ii_* and *γ_ii_*, are not penalized since column *m* only has zero values. The penalty matrix in (11) is now reformulated as follows:

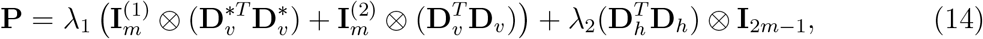

where 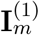 and 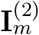 are diagonal indicator matrices given by:

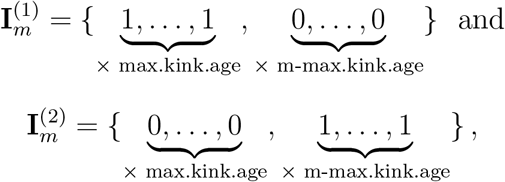

where *max.kink.age* indicates the maximum age at which a kink on the main diagonal is possible. In this paper, we calibrate *max.kink.age* = 31 (i.e., {0,…, 30} years), although a sensitivity analysis with higher values for *max.kink.age* yielded quantitatively similar results. In penalty matrix (12), a similar adjustment is applied to the matrix **D**_*h*_. It is worth noting that social contact rates on the main diagonal that are adjusted by the kink are still penalized in the dimension of the cohort to assure that smooth contact rates are obtained on the diagonals of the contact matrix. The introduction of this kink thus leads to a smoothed contact surface that is non-differentiable on the main diagonal in the dimension of the contact’s age.

### 2.5 Negative Binomial Likelihood

Using the Poisson distribution for the observed contacts *y_ij_* implies that the mean and the variance are equal, i.e. *E*(*Y_ij_*) = Var(*Y_ij_*), while in practice, contact data often display overdispersion. Not accounting for possible overdispersion can lead to biased results. We therefore also impose a negative Binomial distribution for the observed contacts, namely *y_ij_* ~ NegBin(*μ_ij_*, *α_ij_*). The use of a negative Binomial distribution implies that E(*Y_ij_*) = *μ_ij_* and 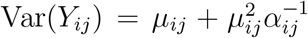. Here, we consider the parameterization with *α_ij_* = *μ_ij_ϕ*^-1^, where *ϕ* > 0 denotes the disperion parameter and the variance is given by Var(*Y_ij_*) = *μ_ij_*(1 + *ϕ*). In the limiting case where *ϕ* tends to zero, the mean and variance will be equal. Note that the variance term resembles the error term of an overdispersed Poisson distribution (Nelder and Lee, 1992). The alternative parameterization with *α_ij_* = *ϕ*^-1^ (leading to Var(*Y_ij_*) = *μ_ij_*(1 + *ϕμ_ij_*)) was also explored but not further described since it performed worse in terms of AIC for the application on the Belgian contact data.

In case *ϕ* is fixed at a certain value, parameter estimates 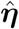 are again obtained through C-PIRLS. The only adaptation is that the entries of **W**^[*k*]^ are given by 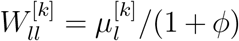. Rather than fixing *ϕ* at a certain value, we are also interested in obtaining a data-driven estimate of *ϕ*. In that endeavor, a two-stage iteration scheme is undertaken, namely by iterating and cycling between holding *ϕ* fixed and holding ***η*** fixed at its current estimate. More specifically, by holding *ϕ* fixed at the current estimate 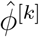, estimates 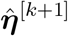 are obtained through C-PIRLS. Next, ***η*** is fixed at 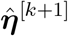 and an updated estimate 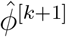 is obtained using the moment estimator (Breslow, 1984). This process is iterated until convergence. Moment estimation of *ϕ* is based on the Pearson’s chi-squared statistic (Breslow, 1984), namely:

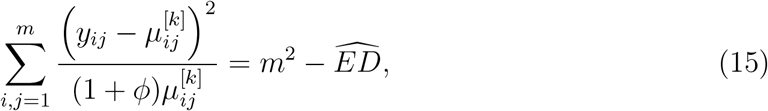

where 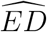 is the trace of the matrix given in (7). This leads to a straightforward estimate of 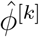:

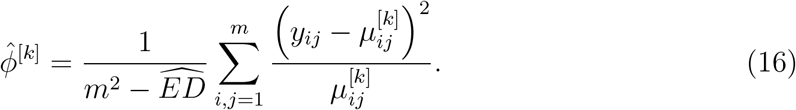

Optimal smoothing parameters λ_1_ and λ_2_ are again chosen via a grid search using the criterion 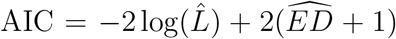. Adding 1 to 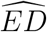 accounts for the estimation of the additional *ϕ* parameter in the negative Binomial setting.

### 2.6 Quantifying the uncertainty of estimates

In order to quantify the uncertainty of the estimate 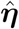, we need to compute its associated variance-covariance matrix. For this purpose, we follow Wood (2006) and use a Bayesian approach to determine the posterior variance-covariance matrix by:

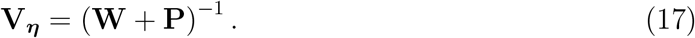

Moreover, as justified by large sample results, the corresponding posterior distribution is taken to be multivariate normal:

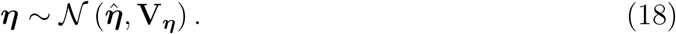

The above (approximate) posterior distribution can be used to calculate confidence intervals for parameters *η_ij_* or for non-linear functions of these parameters (such as *γ_ij_*). An estimate of **V_*η*_** can be obtained by plugging in **W** at convergence together with the estimated optimal smoothing parameters 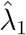 and 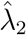 in **P**. The result in (18) can also be used to generate *new* social contact matrices by sampling from the obtained multivariate Gaussian distribution. This can be extremely useful to acknowledge the variability originating from social contact data in the estimation of epidemiological parameters and/or health economic evaluations (Bilcke et al., 2011). Further computational and algorithmic considerations related to C-PIRLS are given in appendix B.

## 3 Simulation Study

A comparison of the methods introduced in Sections 2.1–2.3 is implemented via a simulation study, where the observed contact rates are generated from a Poisson and negative Binomial distribution respectively. We investigate a scenario in which no kink is needed on the main diagonal, and another scenario in which the kink is specified.

Our data generating process is based on a so-called true social contact matrix, denoted by **Γ**\*, from which data is simulated. To obtain such a matrix, a non-parametric regression is applied to the Belgian social contact data. More specifically, the observed contacts rates (see Figure 2 left panel), *y_ij_*/*r_i_* are smoothed using local linear regression. Using a local linear regression approach, there is no guarantee that **K**\* ≡ **Γ**\* ⊙**P** is symmetric. Therefore, we derive a simple symmetric matrix from **K**\*, denoted by 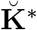, computed as:

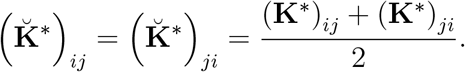

**Figure 2:**
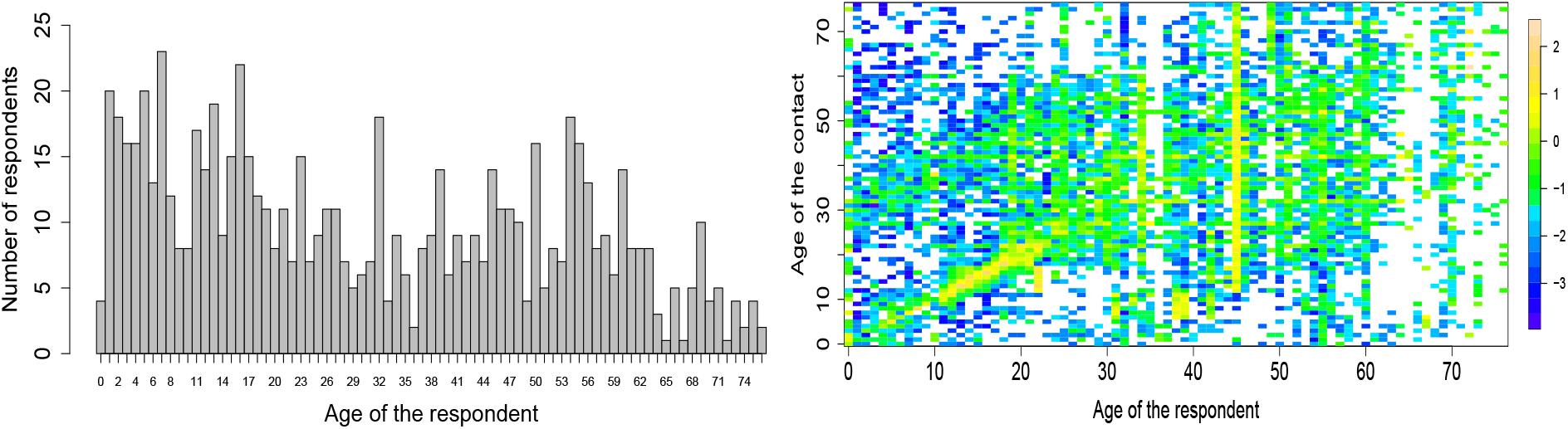
The number of respondent per age (left panel) and the observed log-contact rates (log(*y_ij_*/*r_i_*)) (right panel) of the Belgian social contact data. A white cell indicates that there were no contacts observed for those particular ages of the respondents and contacts.

The true contact surface, 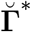 that is used for data simulation is obtained by 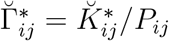. Finally, we denote the log-transformed matrix by 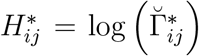. In Figure 3, the true social contact matrices used to generate the data for the simulation study 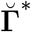 and **H**\* are shown. To account for a kink in the simulation study, we proceed as follows. Let 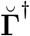 denote the true social contact matrix with a kink on the main diagonal. Matrix 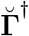 is similar as matrix 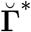, with the exception that the values of 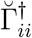, for *i* = 1,…, 24, are artificially increased in the following manner:

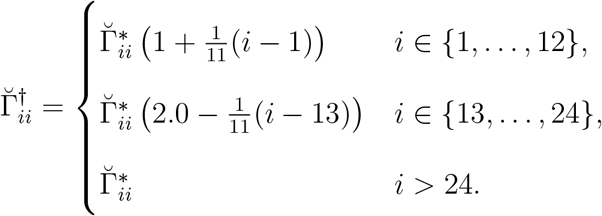

**Figure 3:**
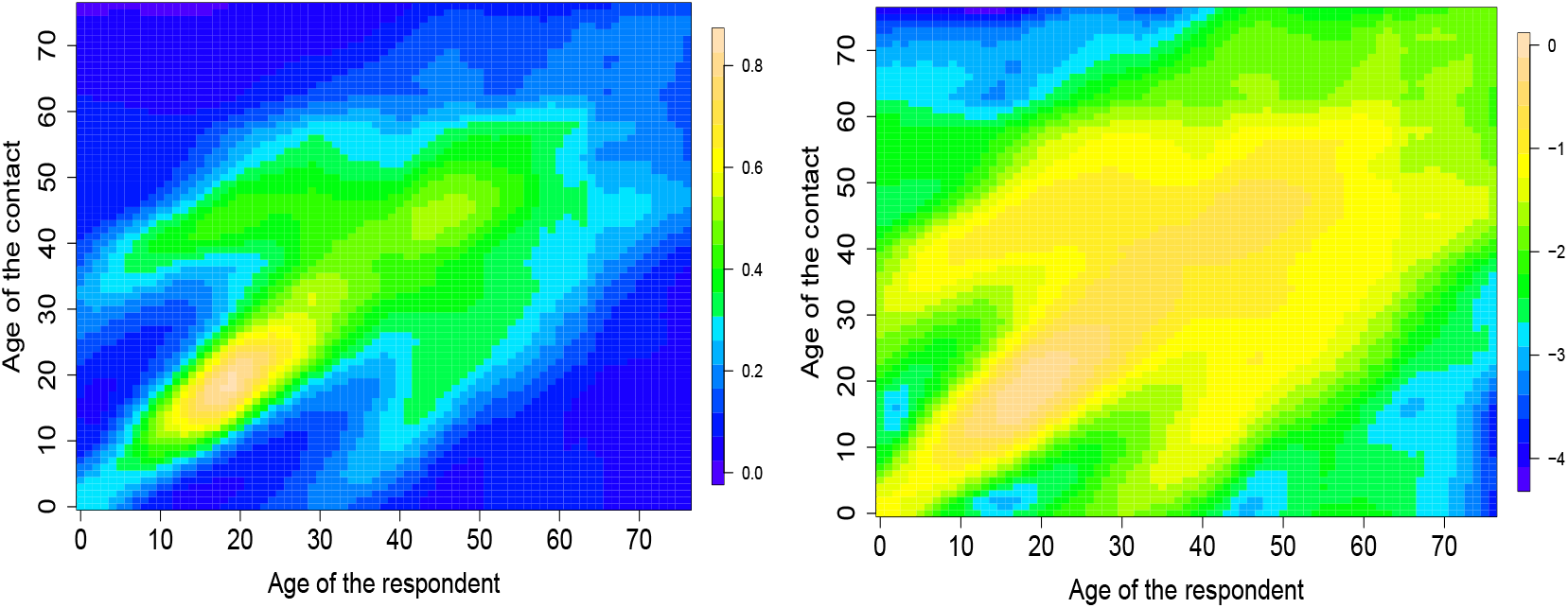
True social contact matrices 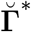 (left) and **H**\* (right) used for the data generating process in the simulation study. The true social contact surfaces are obtained from a nonparametric regression using a local linear fit to the Belgian social contact data.

Thus for ages between 0 and 23 a higher number of contacts is obtained on the main diagonal. Data is simulated using the same participant distribution as in the Belgian social contact data with sample size *n* = 745 (see Figure 2 left). For the Poisson distribution, data is simulated as follows:

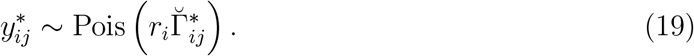

For the negative Binomial distribution (with *ϕ* = 2), the observed number of contacts are obtained as:

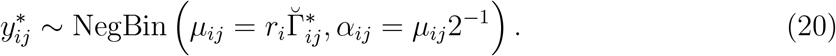

We simulate *S* = 100 datasets for each distributional setting (i.e. Poisson and negative Binomial) and fit models 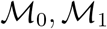 and 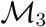 to each dataset with and without consideration of a kink. Optimal smoothing parameters are obtained via grid search using the AIC. This yields estimated social contact matrices 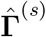 and **Ĥ**^(*s*)^, for *s* = 1,…, *S*. The estimation performance of the different methods are compared using the squared bias and mean square error (MSE). These scalar measures of performance are given by:

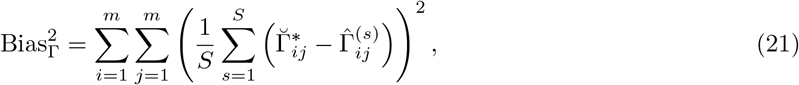

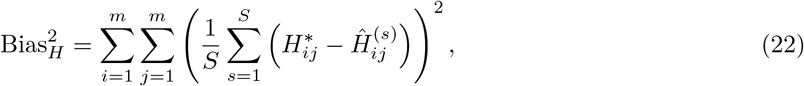

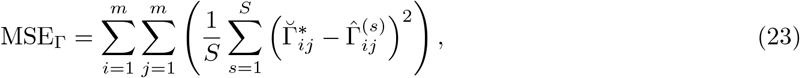

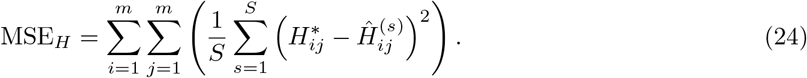

Besides the performance of pointwise estimators given in Tables 1 and 2, we also assess the accuracy with which uncertainty is quantified by looking at the coverage performance of 95% pointwise confidence intervals (CIs) of *η_ij_* in Table 3. Using the approximate posterior distribution in (18), 95% pointwise CIs are easily calculated (i.e., ±1.96× the square root of the Bayesian posterior variance). The reported nominal coverages of the CIs are calculated by averaging over the entries of the entire social contact matrix.

**Table 1:**
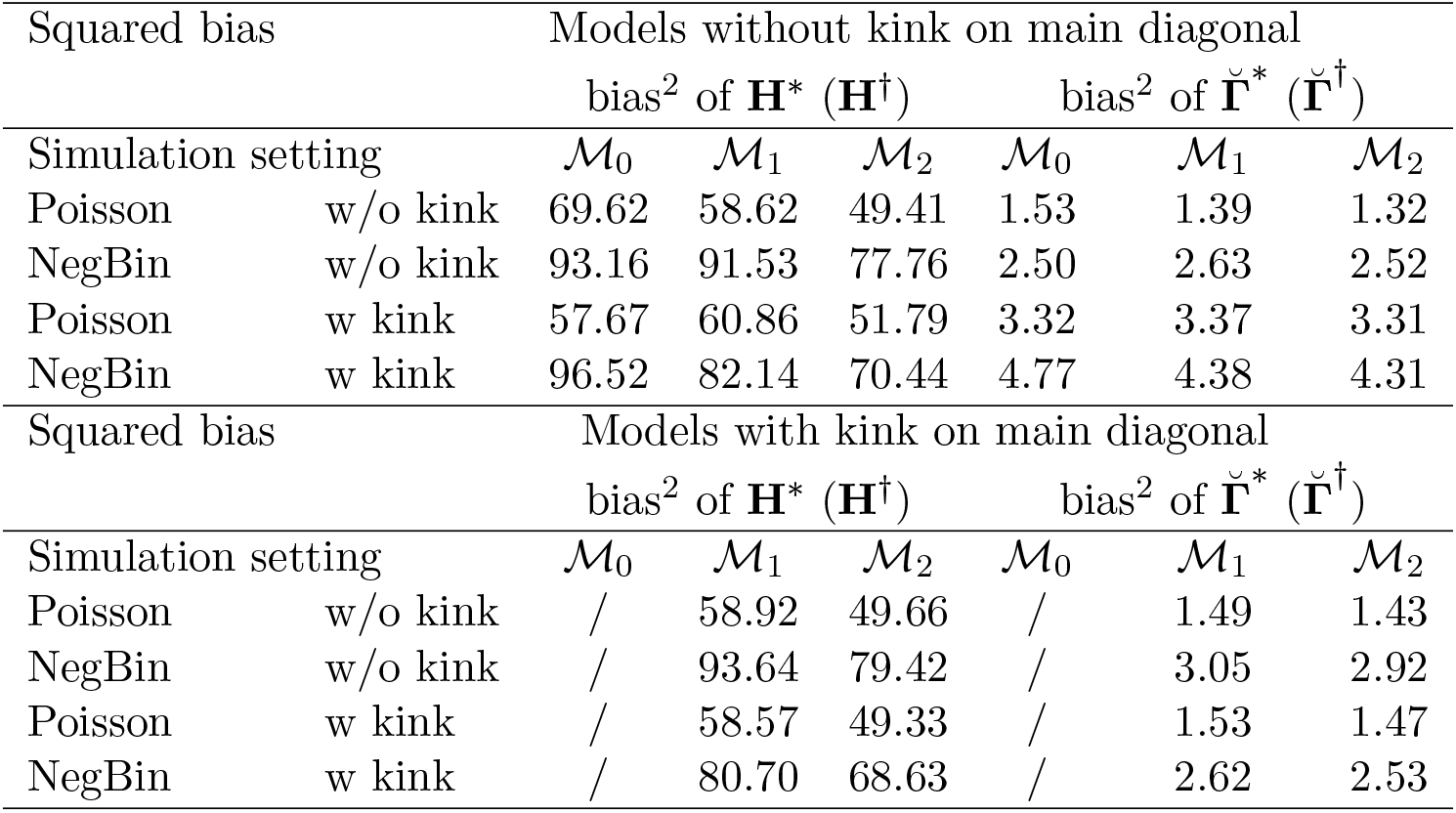
Squared bias of the social contact matrices **H**\* and 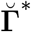 over *S* = 100 simulations using 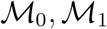 and 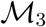 with and without a kink on the main diagonal.

| Squared bias |  | Models without kink on main diagonal |  |  |  |  |  |
| --- | --- | --- | --- | --- | --- | --- | --- |
| | | bias<sup>2</sup> of $\mathbf{H}^*$ ($\mathbf{H}^\dagger$) | | | bias<sup>2</sup> of $\check{\mathbf{\Gamma}}^*$ ($\check{\mathbf{\Gamma}}^\dagger$) | | |
| Simulation setting | | $\mathcal{M}_0$ | $\mathcal{M}_1$ | $\mathcal{M}_2$ | $\mathcal{M}_0$ | $\mathcal{M}_1$ | $\mathcal{M}_2$ |
| Poisson | w/o kink | 69.62 | 58.62 | 49.41 | 1.53 | 1.39 | 1.32 |
| NegBin | w/o kink | 93.16 | 91.53 | 77.76 | 2.50 | 2.63 | 2.52 |
| Poisson | w kink | 57.67 | 60.86 | 51.79 | 3.32 | 3.37 | 3.31 |
| NegBin | w kink | 96.52 | 82.14 | 70.44 | 4.77 | 4.38 | 4.31 |
| Squared bias |  | Models with kink on main diagonal |  |  |  |  |  |
| | | bias<sup>2</sup> of $\mathbf{H}^*$ ($\mathbf{H}^\dagger$) | | | bias<sup>2</sup> of $\check{\mathbf{\Gamma}}^*$ ($\check{\mathbf{\Gamma}}^\dagger$) | | |
| Simulation setting | | $\mathcal{M}_0$ | $\mathcal{M}_1$ | $\mathcal{M}_2$ | $\mathcal{M}_0$ | $\mathcal{M}_1$ | $\mathcal{M}_2$ |
| Poisson | w/o kink | / | 58.92 | 49.66 | / | 1.49 | 1.43 |
| NegBin | w/o kink | / | 93.64 | 79.42 | / | 3.05 | 2.92 |
| Poisson | w kink | / | 58.57 | 49.33 | / | 1.53 | 1.47 |
| NegBin | w kink | / | 80.70 | 68.63 | / | 2.62 | 2.53 |

**Table 2:**
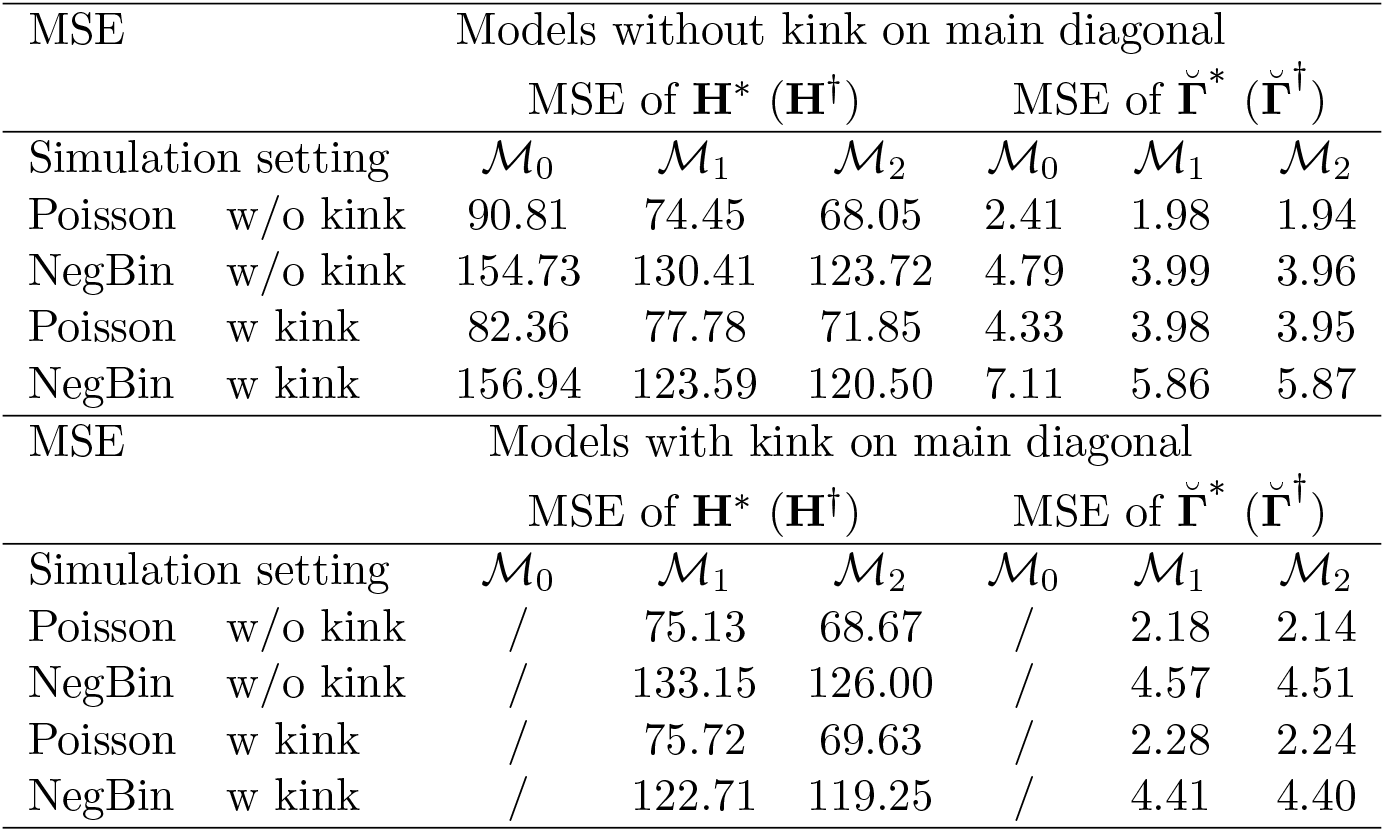
Mean square error of the social contact matrices **H**\* and 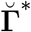 over *S* = 100 simulations using 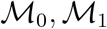 and 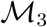 with and without a kink on the main diagonal.

**Table 3:**
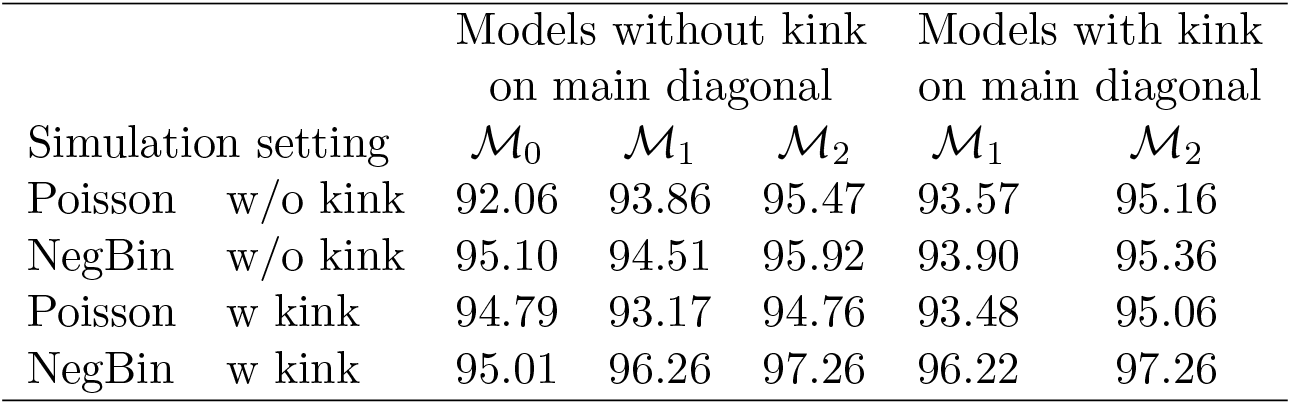
Nominal coverage of 95% pointwise confidence intervals of the social contact matrices **H**\* (**H**^†^) over *S* = 100 simulations using 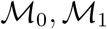 and 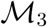 with and without a kink on the main diagonal. The nominal coverage is calculated by averaging over the entire social contact matrix.

For all simulation settings, we observe that models that smooth over cohorts (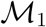 and 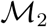) are performing better in terms of MSE than 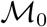, and this holds for both **H**\* and 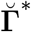. In terms of bias, the results are less clear, but overall model 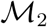 is performing best. When comparing models 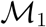 and 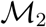, we observe that the latter model has better performance. In the simulation settings in which no kink is introduced on the main diagonal, we observe that models with a kink on the main diagonal perform slightly worse than those without a kink. However, in the simulation settings with a kink, a more pronounced difference is observed in favour of the models with a kink on the main diagonal, especially for 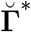. The better performance of models including a kink is mainly due to the better estimation of the main diagonal components of the social contact matrix. No meaningful differences are observed outside the main diagonal region.

In the negative Binomial simulation setting, the overdisperion parameter *ϕ* is estimated well. In the simulation setting without a kink, model 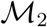 without a kink has an average estimate for *ϕ* of 1.92 with 95% of the estimated overdispersion parameters between 1.74 and 2.22. For the simulation setting with a kink, we find 1.93 (1.71 - 2.20) for model 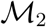.

Table 3 highlights the nominal coverage results of the different simulation settings. We observe that all methods produce pointwise CIs with close to 95% nominal coverage. In the last simulation setting (the negative Binomial distribution with a kink on the main diagonal), a slight overcoverage is observed for models 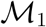 and 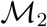. In the latter scenarios, the average lengths of the 95% CIs are 0.65, 0.61 and 0.60, for 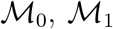 and 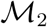 with a kink, respectively. This implies that the overcoverage is not directly associated with wider CIs. Finally, the results in Table 3 indicate that the large sample result in (18) can be used to construct CIs with appropriate nominal coverage.

## 4 Application: Belgian Social Contact Data

The proposed smoothing methods are illustrated on the POLYMOD social contact data of Belgium, obtained through a population-based contact survey carried out over the period of March to May 2006. Participants kept a paper diary with information on their contacts over one day. A contact was defined as a two-way conversation of at least three words in each other’s proximity and the gathered information included the age of the contact, gender, location, duration, frequency, and whether or not touching was involved. Sampling weights – the inverse of the probability that an observation is included because of the sampling design – are available for each participant, based on official age and household size data of the year 2000 census published by Eurostat (Mossong et al., 2008). To estimate population-related social contacts, these sampling weights are included in the analysis. We consider the contact data of all participants aged between 0 and 76 years (both included). In total, we have information on 745 participants from which 399 (53.6%) are females and 345 (46.3%) are males (the information on gender was omitted for one participant). The mean age of the respondents is 31 years. We also restrict to contacts made with individuals between 0 and 76 years (both included), resulting in a total of 13 493 contacts. This gives a crude mean of 18.1 contacts per participant. Furthermore, the age structure of the general population in which the contact survey is conducted in 2006 is obtained from Eurostat (Eurostat, 2017), where the population size in the 0-76 years interval is *N*=9 777 488.

In Figure 2 (right panel), the observed log-contact rates log(*y_ij_*/*r_i_*) of the POLYMOD Belgian social contact data are shown. To estimate the social contact rates from these data, we use models 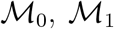 and 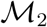 under a Poisson and negative Binomial assumption on the number of contacts with and without a kink on the main diagonal. Let 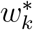 denote the normalized sampling weight of respondent *k*, with *k* = 1,…, 745. The *ij*th input of **Y** is constructed as 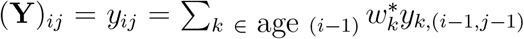 and corresponds to a weighted sum of the number of contacts made by respondents of age *i* – 1 with contacts of age *j* – 1. It follows that the inputs of the vector **r** are given by 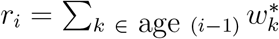.

In Table 4, summary results of the fitted models are given. It can be seen that the negative Binomial distribution performs better in terms of the AIC (smaller AIC is “better”), so that the assumption of a variance that is linearly dependent on the mean is preferred. The effective degrees of freedom 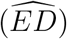 for the Poisson case are higher, indicating that the Poisson distribution tries to explain the observed variability through the mean. From here, we focus on the results of the negative Binomial model. It can be observed that approaches 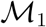 and 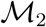 including a kink are performing better in terms of the AIC as compared to those without a kink. Regarding the estimated smoothing parameters 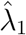 and 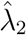, an interesting difference is observed between 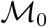 and models 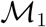 and 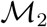. In 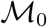, the optimal values for 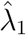 and 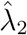 are similar, while for the models accounting for cohort smoothing, the optimal value for 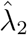 is larger than 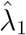, indicating that more penalization is needed in the direction of the cohorts. In terms of computational speed, fitting model 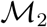 is much faster (≈ 4 times faster) as compared to model 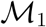, as for the latter model 2*m*^2^ – *m* = 11 781 parameters (including *m*^2^ – *m* nuisance parameters) need to be estimated, as compared to *m*^2^ = 5 929 parameters for 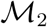.

**Table 4:** Summary results of the fitted models to the Belgian social contact data. Estimated smoothing parameters, effective degrees of freedom, −2 times log-likelihood, AIC and *ϕ* are provided.

| Model | $\hat{\lambda}_1$ | $\hat{\lambda}_2$ | $\widehat{ED}$ | $-2 \log(\hat{L})$ | AIC | $\hat{\phi}$ |
| --- | --- | --- | --- | --- | --- | --- |
| $\mathcal{M}_0$ Poisson | 0.46 | 0.46 | 1 979.5 | 20 151.7 | 24 110.7 | - |
| $\mathcal{M}_0$ NegBin | 15.17 | 16.64 | 181.5 | 20 732.7 | 21 097.7 | 3.08 |
| $\mathcal{M}_1$ Poisson | 0.50 | 0.32 | 2 085.6 | 20 318.4 | 24 489.6 | - |
| $\mathcal{M}_1$ NegBin | 22.76 | 1714.91 | 55.8 | 20 988.5 | 21 102.1 | 3.76 |
| $\mathcal{M}_2$ Poisson | 0.50 | 0.32 | 2 125.0 | 20 287.6 | 24 537.7 | - |
| $\mathcal{M}_2$ NegBin | 27.36 | 1564.02 | 59.5 | 20 994.2 | 21 115.2 | 3.76 |
| With kink |  |  |  |  |  |  |
| $\mathcal{M}_1$ NegBin | 30.00 | 1584.89 | 54.3 | 20 967.6 | 21 078.2 | 3.70 |
| $\mathcal{M}_2$ NegBin | 40.00 | 1584.89 | 55.4 | 20 973.2 | 21 086.0 | 3.70 |

In Figures 4 and 5, the estimated log contact rate surfaces, **Ĥ**, and the mixing at the population level, 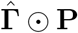, for models 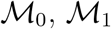 and 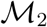 under the negative Binomial distribution without a kink are shown. Generally, the surfaces are able to capture important features of human contact behaviour. There is a clear difference in the estimated surfaces for model 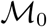 and models 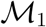 and 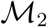 in the sense that diagonal components are more pronounced for models accounting for cohort smoothing. The shifted diagonal between children and parents is also more clearly visible.

**Figure 4:**
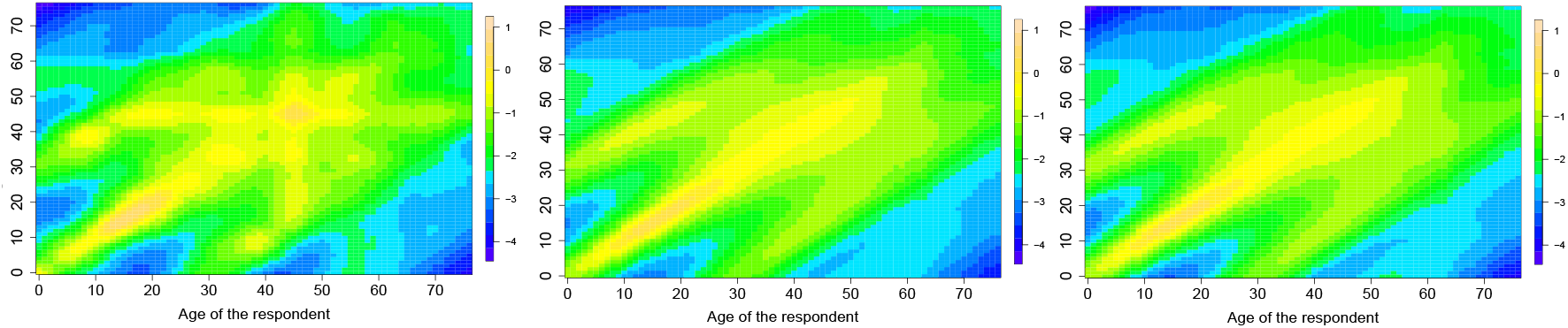
The estimated log contact rates surface, **Ĥ**, for models 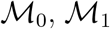 and 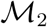 without kink (left to right) with the negative Binomial distribution.

**Figure 5:**
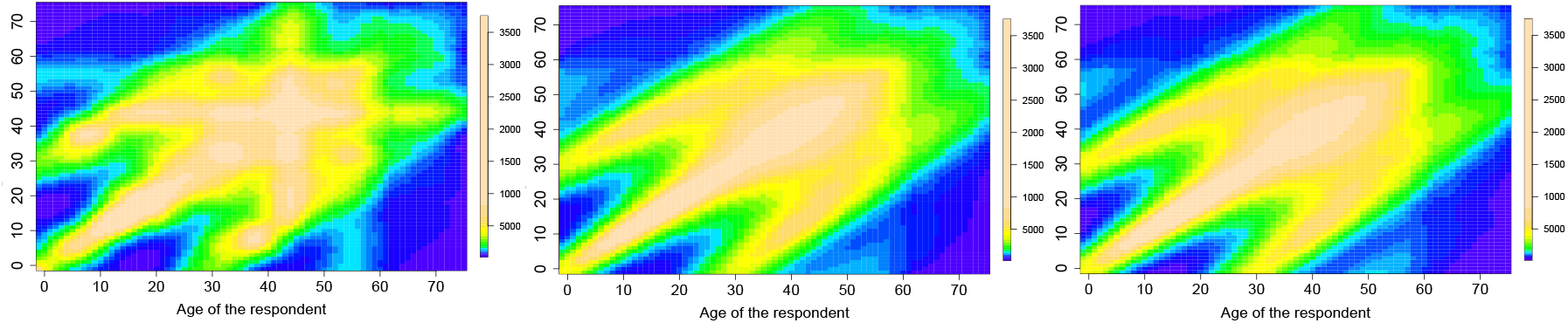
The estimated mixing at the population level, 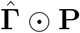, for models 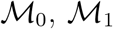 and 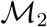 without kink (left to right) with the negative Binomial distribution.

Based on the AIC values in Table 4, we see that the models including a kink are preferred. In addition, based on the results of the simulation study in Section 3, the estimated contact rates are very similar for models 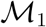 and 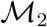, so that we prefer the use of model 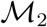 for the POLYMOD Belgian social contact data as it is less computationally intensive.

In Figure 6, estimated contact surfaces are shown for model 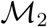 with the negative Binomial distribution and a kink on the main diagonal. From the figure on the bottom, it is observed that the main diagonal has higher values for younger ages for the model including the kink and this yields higher values on the main diagonal of **Ĥ** and 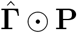. For the model where the kink is absent, the values in the estimated matrix 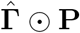 range from 1 496.2 to 162 986.5, whereas for the model with kink, the values range from 1 608.4 to 375 371.5. The kink thus allows for a huge increase in the estimated number of contacts for children and young adults with individuals of the same age. These results enforce the fact that a kink is needed to capture the non-smooth effect of mixing with people of the same age, especially for the children and young adults.

**Figure 6:**
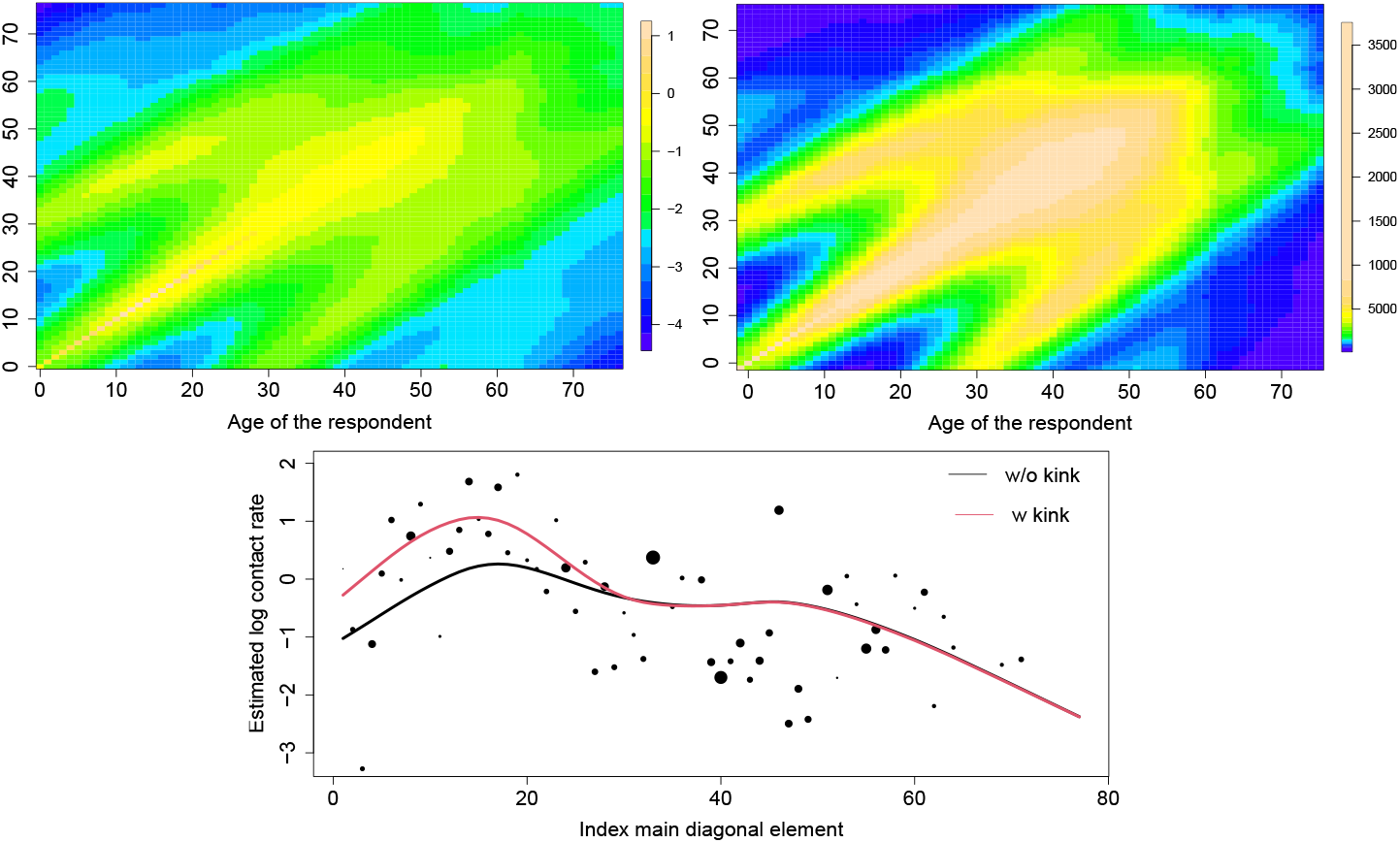
The estimated log contact rates surface (left), **Ĥ**, and the mixing at the population level (right), 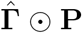, for model 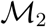 with the negative Binomial distribution including an additional kink on the main diagonal. The diagonal elements of **Ĥ** for the model with and without a kink (bottom), together with the observed log-contact rates.

## 5 Discussion

Quantifying contact behaviour contributes to a better understanding of how infectious diseases spread (Anderson and May, 1991; Edmunds et al., 1997). Social contact rates play a major role in mathematical models used to model infectious disease transmission. In this paper, we describe a smoothing constrained approach to estimate social contact rates from self-reported social contact data. The proposed approach assumes that the contact rates are smooth from a cohort perspective as well as from the age distribution of contacts. Thus, besides smoothing in the direction of the age of contacts, we propose to smooth contact rates from a cohort perspective by following two alternative strategies.

The simulation study and the data application show that approach 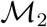, in which the penalty matrix is reordered (and penalization is performed over the diagonal components), is performing better. It was observed that this method yielded the smallest MSE over all simulation settings. Additionally, confidence intervals with nominal coverage close to 95% were obtained. In the Belgian data application, the computation time of method 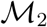 is three to four times faster than method 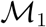, and so we recommend the use of the former approach for the estimation of social contact rates. The true social contact surface used in the data generating process of the simuation study was obtained through local linear regression of the raw social contact rates of the Belgian POLYMOD study. This approach is preferred for two reasons. First, by using the same data in the simulation study as in the application presented in Section 4, a better view of the performance of the different approaches can be obtained. Second, we are not aware of any easy applicable mathematical formula or fully parametric model of a two dimensional surface that would be suitable to represent a contact rate surface. A search is needed to calibrate the smoothing parameters λ_1_ and λ_2_. This is a disadvantage compared to the approach by van de Kassteele et al. (2017) in which the amount of smoothing is directly estimated together with model parameters from the information in the data. However, with the availability of fast parallel computing and multi-core machines, the grid search can be performed relatively fast.

In this paper, the contact rates are assumed indifferent for men and women. Recently, van de Kassteele et al. (2017) presented a Bayesian model for estimating social contact rates for men and women, with results suggesting that different contact patterns exist and thus that there is a gender effect. Future work could investigate how the methodology proposed in this paper can be extended to estimate social contact rates between both sexes without increasing the computational burden. A comparison with other methods used to smooth social contact data was not done in this paper. Future extensions could focus on the impact of social contact matrices obtained from different methods on key epidemiological parameters. In general, age-specific contact rates are also used as an input in the comparison and evaluation of vaccination schedules via future projections (Beutels et al., 2013). Most evaluations assume a fixed social contact rate matrix and thus that no uncertainty is related to this input. The result derived in equation (18) offers a tool to account for the variability associated with the estimation of social contact rates. By simulation of new contact matrices from (18), the associated variability can be taken into account in the evaluation of vaccination strategies and related health economic evaluations.

Finally, our proposed methodology does not employ any regression basis such as B-splines because an exact link between the constraints and linear predictors is needed. We are exploring whether the proposed methodology can be extended to make use of basis functions that will likely lead to a reduction of the computational cost. Alternative ways of incorporating the reciprocal nature of the phenomenon will thus be necessary.

## Acknowledgments

For the simulation study with the negative Binomial model, we used the infrastructure of the VSC–Flemish Supercomputer Center, funded by the Hercules Foundation and the Flemish Government– department EWI. Support from the University of Antwerp scientific chair in Evidence-Based Vaccinology, financed in 2009-2017 by a gift from Pfizer, is acknowledged [to NH]. This project has received funding from the European Research Council (ERC) under the European Union’s Horizon 2020 research and innovation programme (grant agreement 682540 - TransMID). Oswaldo Gressani, Niel Hens and Christel Faes would also like to thank the European Union’s Research and Innovation Action EpiPose (grant number 101003688) for funding this work.

## Data availability statement

Code to reproduce the results of this paper is available at https://github.com/oswaldogressani/Cohort_smoothing.

## Appendix

## Appendix A. Penalty matrix P_*d*_

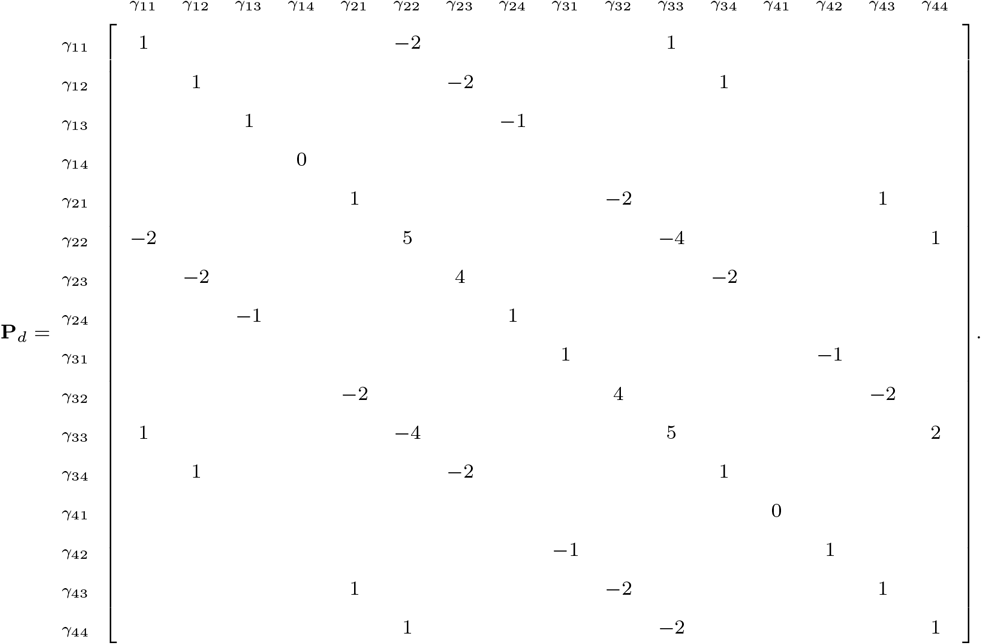

## Appendix B. Computational considerations

`R` version 4.1.2 is used to fit the proposed models. To enhance convergence of the proposed C-PIRLS fitting scheme, we first perform parameter estimation using penalized iterative reweighted least squares without using the symmetry constraint and use the obtained estimated parameters as starting values in the C-PIRLS algorithm. To initiate the estimation of PIRLS without the symmetry constraint, starting values 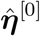 are needed. These can, for example, be set at 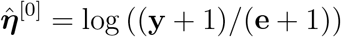.

The same number of parameters are estimated as there are entries in the matrices **Γ** or 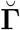. For instance, in our application (*m* = 77) in Section 4, we need to estimate *m*^2^ = 5 929 and 2*m*^2^ – *m* = 11 781 parameters, respectively. This is practically challenging on a regular personal computer. Therefore, we make use of sparse matrix implementations by using the `R`-package `Matrix` (Bates and Maechler, 2017).

To choose the optimal smoothing parameters λ_1_ and λ_2_, a grid search is performed with both λ_1_,λ_2_ ∈ {0.5, 1, 5, 10, 50, 100, 500, 1000, 5000, 10000}. This initial grid search gives an indication (based on minimization of the AIC) of the values of the optimal smoothing parameters. In a second step, a greedy grid search is performed an a denser grid using the `cleversearch` function in the `R`-package `svcm` (Heim, 2007).

