## Supplementary Materials for "Cohort-based smoothing methods for age-specific contact rates"

Last update: April 4, 2022

### Examples of the defined vectors and matrices in Section 2

Here, we present an overview of the defined vectors and matrices that are defined throughout Section 2 of the manuscript for an example with $m = 4$. The vector $\mathbf{r} = (r_i)$ containing the total number of respondents of age $i - 1$, and the vector $\mathbf{p} = (p_i)$ containing the population size of individuals of age $i - 1$ are:

$$\mathbf{r} = \begin{pmatrix} r_1 \\ r_2 \\ r_3 \\ r_4 \end{pmatrix} \quad \text{and} \quad \mathbf{p} = \begin{pmatrix} p_1 \\ p_2 \\ p_3 \\ p_4 \end{pmatrix}.$$

The matrix $\mathbf{Y}$ containing the total number of contacts made by the respondents of age $i - 1$ with individuals of age $j - 1$, the matrix $\mathbf{E} = \mathbf{r}\mathbf{1}_m$ and the $\mathbf{P} = \mathbf{p}\mathbf{1}_m$ are:

$$\mathbf{Y} = \begin{pmatrix} y_{11} & y_{12} & y_{13} & y_{14} \\ y_{21} & y_{22} & y_{23} & y_{24} \\ y_{31} & y_{32} & y_{33} & y_{34} \\ y_{41} & y_{42} & y_{43} & y_{44} \end{pmatrix}, \mathbf{E} = \begin{pmatrix} r_1 & r_1 & r_1 & r_1 \\ r_2 & r_2 & r_2 & r_2 \\ r_3 & r_3 & r_3 & r_3 \\ r_4 & r_4 & r_4 & r_4 \end{pmatrix} \quad \text{and} \quad \mathbf{P} = \begin{pmatrix} p_1 & p_1 & p_1 & p_1 \\ p_2 & p_2 & p_2 & p_2 \\ p_3 & p_3 & p_3 & p_3 \\ p_4 & p_4 & p_4 & p_4 \end{pmatrix}.$$

The vectors $\mathbf{y}$ and $\mathbf{e}$ obtained by arranging the matrices $\mathbf{Y}$ and $\mathbf{E}$, respectively, in row order into a vector, are:

$$\mathbf{y} = \begin{pmatrix} y_{11} \\ y_{12} \\ y_{13} \\ y_{14} \\ y_{21} \\ y_{22} \\ y_{23} \\ y_{24} \\ y_{31} \\ y_{32} \\ y_{33} \\ y_{34} \\ y_{41} \\ y_{42} \\ y_{43} \\ y_{44} \end{pmatrix} \quad \text{and} \quad \mathbf{e} = \begin{pmatrix} r_1 \\ r_1 \\ r_1 \\ r_1 \\ r_2 \\ r_2 \\ r_2 \\ r_2 \\ r_3 \\ r_3 \\ r_3 \\ r_3 \\ r_4 \\ r_4 \\ r_4 \\ r_4 \end{pmatrix}.$$

The social contact matrix $\mathbf{\Gamma}$ containing the actual contact rates is:

$$\mathbf{\Gamma} = \begin{pmatrix} \gamma_{11} & \gamma_{12} & \gamma_{13} & \gamma_{14} \\ \gamma_{21} & \gamma_{22} & \gamma_{23} & \gamma_{24} \\ \gamma_{31} & \gamma_{32} & \gamma_{33} & \gamma_{34} \\ \gamma_{41} & \gamma_{42} & \gamma_{43} & \gamma_{44} \end{pmatrix}.$$

The vector $\boldsymbol{\gamma}$ obtained by arranging the matrix $\boldsymbol{\gamma}$ by row order into a vector, the vector $\boldsymbol{\mu} = \mathbf{e} \odot \boldsymbol{\gamma}$, and the vector $\boldsymbol{\eta} = \log(\boldsymbol{\gamma})$ are:

$$
\boldsymbol{\gamma} = \begin{pmatrix} \gamma_{11} \\ \gamma_{12} \\ \gamma_{13} \\ \gamma_{14} \\ \gamma_{21} \\ \gamma_{22} \\ \gamma_{23} \\ \gamma_{24} \\ \gamma_{31} \\ \gamma_{32} \\ \gamma_{33} \\ \gamma_{34} \\ \gamma_{41} \\ \gamma_{42} \\ \gamma_{43} \\ \gamma_{44} \end{pmatrix}, \boldsymbol{\mu} = \begin{pmatrix} \mu_{11} \\ \mu_{12} \\ \mu_{13} \\ \mu_{14} \\ \mu_{21} \\ \mu_{22} \\ \mu_{23} \\ \mu_{24} \\ \mu_{31} \\ \mu_{32} \\ \mu_{33} \\ \mu_{34} \\ \mu_{41} \\ \mu_{42} \\ \mu_{43} \\ \mu_{44} \end{pmatrix} = \begin{pmatrix} r_1\gamma_{11} \\ r_1\gamma_{12} \\ r_1\gamma_{13} \\ r_1\gamma_{14} \\ r_2\gamma_{21} \\ r_2\gamma_{22} \\ r_2\gamma_{23} \\ r_2\gamma_{24} \\ r_3\gamma_{31} \\ r_3\gamma_{32} \\ r_3\gamma_{33} \\ r_3\gamma_{34} \\ r_4\gamma_{41} \\ r_4\gamma_{42} \\ r_4\gamma_{43} \\ r_4\gamma_{44} \end{pmatrix} \quad \text{and} \quad \boldsymbol{\eta} = \begin{pmatrix} \eta_{11} \\ \eta_{12} \\ \eta_{13} \\ \eta_{14} \\ \eta_{21} \\ \eta_{22} \\ \eta_{23} \\ \eta_{24} \\ \eta_{31} \\ \eta_{32} \\ \eta_{33} \\ \eta_{34} \\ \eta_{41} \\ \eta_{42} \\ \eta_{43} \\ \eta_{44} \end{pmatrix}.
$$

The allocation matrix $\boldsymbol{L}$ and the vector $\boldsymbol{\nu}$ of the reciprocal constraint in (2) are:

$$
\mathbf{L} = \begin{array}{c} \begin{array}{cccccccccccccccc} \scriptstyle\gamma_{11} & \scriptstyle\gamma_{12} & \scriptstyle\gamma_{13} & \scriptstyle\gamma_{14} & \scriptstyle\gamma_{21} & \scriptstyle\gamma_{22} & \scriptstyle\gamma_{23} & \scriptstyle\gamma_{24} & \scriptstyle\gamma_{31} & \scriptstyle\gamma_{32} & \scriptstyle\gamma_{33} & \scriptstyle\gamma_{34} & \scriptstyle\gamma_{41} & \scriptstyle\gamma_{42} & \scriptstyle\gamma_{43} & \scriptstyle\gamma_{44} \end{array} \\ \begin{pmatrix} & 1 & & & -1 & & & & & & & & & & & \\ & & 1 & & & & & & -1 & & & & & & & \\ & & & 1 & & & & & & & & & -1 & & & \\ & & & & & & 1 & & & -1 & & & & & & \\ & & & & & & & 1 & & & & & & -1 & & \\ & & & & & & & & & & & 1 & & & -1 & \end{pmatrix} \end{array}
$$

and

$$
\boldsymbol{\nu} = \begin{pmatrix} \log(p_2) - \log(p_1) \\ \log(p_3) - \log(p_1) \\ \log(p_4) - \log(p_1) \\ \log(p_3) - \log(p_2) \\ \log(p_4) - \log(p_2) \\ \log(p_4) - \log(p_3) \end{pmatrix}.
$$

In the penalty matrix (5), the matrices are:

$$\mathbf{D}_h^T\mathbf{D}_h = \mathbf{D}_v^T\mathbf{D}_v = \begin{pmatrix} 1 & -2 & 1 & 0 \\ -2 & 5 & -4 & 1 \\ 1 & -4 & 5 & -2 \\ 0 & 1 & -2 & 1 \end{pmatrix},$$

and thus:

$$\mathbf{I}_4 \otimes (\mathbf{D}_h^T\mathbf{D}_h) = \begin{array}{c} \\ \gamma_{11} \\ \gamma_{12} \\ \gamma_{13} \\ \gamma_{14} \\ \gamma_{21} \\ \gamma_{22} \\ \gamma_{23} \\ \gamma_{24} \\ \gamma_{31} \\ \gamma_{32} \\ \gamma_{33} \\ \gamma_{34} \\ \gamma_{41} \\ \gamma_{42} \\ \gamma_{43} \\ \gamma_{44} \end{array} \begin{array}{c} \begin{array}{cccccccccccccccc} \gamma_{11} & \gamma_{12} & \gamma_{13} & \gamma_{14} & \gamma_{21} & \gamma_{22} & \gamma_{23} & \gamma_{24} & \gamma_{31} & \gamma_{32} & \gamma_{33} & \gamma_{34} & \gamma_{41} & \gamma_{42} & \gamma_{43} & \gamma_{44} \end{array} \\ \left(\begin{array}{cccccccccccccccc}
1 & -2 & 1 & & & & & & & & & & & & & \\
-2 & 5 & -4 & 1 & & & & & & & & & & & & \\
1 & -4 & 5 & -2 & & & & & & & & & & & & \\
 & 1 & -2 & 1 & & & & & & & & & & & & \\
 & & & & 1 & -2 & 1 & & & & & & & & & \\
 & & & & -2 & 5 & -4 & 1 & & & & & & & & \\
 & & & & 1 & -4 & 5 & -2 & & & & & & & & \\
 & & & & & 1 & -2 & 1 & & & & & & & & \\
 & & & & & & & & 1 & -2 & 1 & & & & & \\
 & & & & & & & & -2 & 5 & -4 & 1 & & & & \\
 & & & & & & & & 1 & -4 & 5 & -2 & & & & \\
 & & & & & & & & & 1 & -2 & 1 & & & & \\
 & & & & & & & & & & & & 1 & -2 & 1 & \\
 & & & & & & & & & & & & -2 & 5 & -4 & 1 \\
 & & & & & & & & & & & & 1 & -4 & 5 & -2 \\
 & & & & & & & & & & & & & 1 & -2 & 1
\end{array}\right) \end{array}$$

and

$$
(\mathbf{D}_v^T\mathbf{D}_v)\otimes\mathbf{I}_4 =
\begin{array}{c}
 \\ \gamma_{11} \\ \gamma_{12} \\ \gamma_{13} \\ \gamma_{14} \\ \gamma_{21} \\ \gamma_{22} \\ \gamma_{23} \\ \gamma_{24} \\ \gamma_{31} \\ \gamma_{32} \\ \gamma_{33} \\ \gamma_{34} \\ \gamma_{41} \\ \gamma_{42} \\ \gamma_{43} \\ \gamma_{44}
\end{array}
\begin{array}{c}
\begin{array}{cccccccccccccccc}
\gamma_{11} & \gamma_{12} & \gamma_{13} & \gamma_{14} & \gamma_{21} & \gamma_{22} & \gamma_{23} & \gamma_{24} & \gamma_{31} & \gamma_{32} & \gamma_{33} & \gamma_{34} & \gamma_{41} & \gamma_{42} & \gamma_{43} & \gamma_{44}
\end{array} \\
\begin{pmatrix}
1 & & & & -2 & & & & 1 & & & & & & & \\
 & 1 & & & & -2 & & & & 1 & & & & & & \\
 & & 1 & & & & -2 & & & & 1 & & & & & \\
 & & & 1 & & & & -2 & & & & 1 & & & & \\
-2 & & & & 5 & & & & -4 & & & & 1 & & & \\
 & -2 & & & & 5 & & & & -4 & & & & 1 & & \\
 & & -2 & & & & 5 & & & & -4 & & & & 1 & \\
 & & & -2 & & & & 5 & & & & -4 & & & & 1 \\
1 & & & & -4 & & & & 5 & & & & -2 & & & \\
 & 1 & & & & -4 & & & & 5 & & & & -2 & & \\
 & & 1 & & & & -4 & & & & 5 & & & & -2 & \\
 & & & 1 & & & & -4 & & & & 5 & & & & -2 \\
 & & & & 1 & & & & -2 & & & & 1 & & & \\
 & & & & & 1 & & & & -2 & & & & 1 & & \\
 & & & & & & 1 & & & & -2 & & & & 1 & \\
 & & & & & & & 1 & & & & -2 & & & & 1
\end{pmatrix}
\end{array}.
$$

In Section 2.2 of the manuscript, the newly defined matrices $\breve{\mathbf{\Gamma}}$, $\breve{\mathbf{Y}}$, $\breve{\mathbf{E}}$, $\breve{\mathbf{W}}$ are:

$$
\breve{\mathbf{\Gamma}} = \begin{pmatrix}
\cdot & \cdot & \cdot & \gamma_{41} \\
\cdot & \cdot & \gamma_{31} & \gamma_{42} \\
\cdot & \gamma_{21} & \gamma_{32} & \gamma_{43} \\
\gamma_{11} & \gamma_{22} & \gamma_{33} & \gamma_{44} \\
\gamma_{12} & \gamma_{23} & \gamma_{34} & \cdot \\
\gamma_{13} & \gamma_{24} & \cdot & \cdot \\
\gamma_{14} & \cdot & \cdot & \cdot
\end{pmatrix}, \breve{\mathbf{Y}} = \begin{pmatrix}
\cdot & \cdot & \cdot & y_{41} \\
\cdot & \cdot & y_{31} & y_{42} \\
\cdot & y_{21} & y_{32} & y_{43} \\
y_{11} & y_{22} & y_{33} & y_{44} \\
y_{12} & y_{23} & y_{34} & \cdot \\
y_{13} & y_{24} & \cdot & \cdot \\
y_{14} & \cdot & \cdot & \cdot
\end{pmatrix},
$$

$$
\breve{\mathbf{E}} = \begin{pmatrix}
. & . & . & r_4 \\
. & . & r_3 & r_4 \\
. & r_2 & r_3 & r_4 \\
r_1 & r_2 & r_3 & r_4 \\
r_1 & r_2 & r_3 & . \\
r_1 & r_2 & . & . \\
r_1 & . & . & .
\end{pmatrix} \quad \text{and} \quad \breve{\mathbf{W}} = \begin{pmatrix}
0 & 0 & 0 & 1 \\
0 & 0 & 1 & 1 \\
0 & 1 & 1 & 1 \\
1 & 1 & 1 & 1 \\
1 & 1 & 1 & 0 \\
1 & 1 & 0 & 0 \\
1 & 0 & 0 & 0
\end{pmatrix}.
$$

We remark once again that the *dots* in $\breve{\mathbf{\Gamma}}$ are actually nuisance parameters which are not of direct interest. However, they must be accounted for in the estimation of those parameters which are of interest. The vectors $\breve{\boldsymbol{\gamma}}$, $\breve{\mathbf{y}}$, $\breve{\mathbf{e}}$, $\breve{\mathbf{w}}$ obtained by arranging the matrices $\breve{\mathbf{\Gamma}}$, $\breve{\mathbf{Y}}$, $\breve{\mathbf{E}}$, $\breve{\mathbf{W}}$ by column order into a vector, the vector $\breve{\boldsymbol{\mu}} = \breve{\mathbf{e}} \odot \breve{\boldsymbol{\gamma}} \odot \breve{\mathbf{w}}$, and the vector $\breve{\boldsymbol{\eta}} = \log(\breve{\boldsymbol{\gamma}})$ are thus given by:

$$\breve{\boldsymbol{\gamma}} = \begin{pmatrix} . \\ . \\ . \\ \gamma_{11} \\ \gamma_{12} \\ \gamma_{13} \\ \gamma_{14} \\ . \\ . \\ \gamma_{21} \\ \gamma_{22} \\ \gamma_{23} \\ \gamma_{24} \\ . \\ . \\ \gamma_{31} \\ \gamma_{32} \\ \gamma_{33} \\ \gamma_{34} \\ . \\ . \\ \gamma_{41} \\ \gamma_{42} \\ \gamma_{43} \\ \gamma_{44} \\ . \\ . \\ . \end{pmatrix}, \breve{\mathbf{y}} = \begin{pmatrix} . \\ . \\ . \\ y_{11} \\ y_{12} \\ y_{13} \\ y_{14} \\ . \\ . \\ y_{21} \\ y_{22} \\ y_{23} \\ y_{24} \\ . \\ . \\ y_{31} \\ y_{32} \\ y_{33} \\ y_{34} \\ . \\ . \\ y_{41} \\ y_{42} \\ y_{43} \\ y_{44} \\ . \\ . \\ . \end{pmatrix}, \breve{\mathbf{e}} = \begin{pmatrix} . \\ . \\ . \\ r_1 \\ r_1 \\ r_1 \\ r_1 \\ . \\ . \\ r_2 \\ r_2 \\ r_2 \\ r_2 \\ . \\ . \\ r_3 \\ r_3 \\ r_3 \\ r_3 \\ . \\ . \\ r_4 \\ r_4 \\ r_4 \\ r_4 \\ . \\ . \\ . \end{pmatrix}, \breve{\mathbf{w}} = \begin{pmatrix} 0 \\ 0 \\ 0 \\ 1 \\ 1 \\ 1 \\ 1 \\ 0 \\ 0 \\ 1 \\ 1 \\ 1 \\ 1 \\ 0 \\ 0 \\ 1 \\ 1 \\ 1 \\ 1 \\ 0 \\ 0 \\ 1 \\ 1 \\ 1 \\ 1 \\ 0 \\ 0 \\ 0 \end{pmatrix}, \breve{\boldsymbol{\mu}} = \begin{pmatrix} . \\ . \\ . \\ \mu_{11} \\ \mu_{12} \\ \mu 13 \\ \mu_{14} \\ . \\ . \\ \mu_{21} \\ \mu_{22} \\ \mu_{23} \\ \mu_{24} \\ . \\ . \\ \mu_{31} \\ \mu_{32} \\ \mu_{33} \\ \mu_{34} \\ . \\ . \\ \mu_{41} \\ \mu_{42} \\ \mu_{43} \\ \mu_{44} \\ . \\ . \\ . \end{pmatrix} = \begin{pmatrix} 0 \\ 0 \\ 0 \\ r_1\gamma_{11} \\ r_1\gamma_{12} \\ r_1\gamma 13 \\ r_1\gamma_{14} \\ 0 \\ 0 \\ r_2\gamma_{21} \\ r_2\gamma_{22} \\ r_2\gamma_{23} \\ r_2\gamma_{24} \\ 0 \\ 0 \\ r_3\gamma_{31} \\ r_3\gamma_{32} \\ r_3\gamma_{33} \\ r_3\gamma_{34} \\ 0 \\ 0 \\ r_4\gamma_{41} \\ r_4\gamma_{42} \\ r_4\gamma_{43} \\ r_4\gamma_{44} \\ 0 \\ 0 \\ 0 \end{pmatrix} \text{ and } \breve{\boldsymbol{\eta}} = \begin{pmatrix} . \\ . \\ . \\ \eta_{11} \\ \eta_{12} \\ \eta_{13} \\ \eta_{14} \\ . \\ . \\ \eta_{21} \\ \eta_{22} \\ \eta_{23} \\ \eta_{24} \\ . \\ . \\ \eta_{31} \\ \eta_{32} \\ \eta_{33} \\ \eta_{34} \\ . \\ . \\ \eta_{41} \\ \eta_{42} \\ \eta_{43} \\ \eta_{44} \\ . \\ . \\ . \end{pmatrix}.$$

The allocation matrix $\mathbf{L}$ in (8) is now:

$$\mathbf{L}^T = \begin{array}{c} \cdot \\ \cdot \\ \cdot \\ \gamma_{11} \\ \gamma_{12} \\ \gamma_{13} \\ \gamma_{14} \\ \cdot \\ \cdot \\ \gamma_{21} \\ \gamma_{22} \\ \gamma_{23} \\ \gamma_{24} \\ \cdot \\ \cdot \\ \gamma_{31} \\ \gamma_{32} \\ \gamma_{33} \\ \gamma_{34} \\ \cdot \\ \cdot \\ \gamma_{41} \\ \gamma_{42} \\ \gamma_{43} \\ \gamma_{44} \\ \cdot \\ \cdot \\ \cdot \end{array} \begin{pmatrix} & & & & & \\ & & & & & \\ & & & & & \\ & & & & & \\ 1 & & & & & \\ & 1 & & & & \\ & & 1 & & & \\ & & & & & \\ & & & & & \\ -1 & & & & & \\ & & & & & \\ & & & 1 & & \\ & & & & 1 & \\ & & & & & \\ & & & & & \\ & -1 & & & & \\ & & & -1 & & \\ & & & & & \\ & & & & & 1 \\ & & & & & \\ & & & & & \\ & & -1 & & & \\ & & & & -1 & \\ & & & & & -1 \\ & & & & & \\ & & & & & \\ & & & & & \\ & & & & & \end{pmatrix}$$

and the vector $\boldsymbol{\nu}$ is similar as defined above. In the penalty matrix (11), the matrix $\mathbf{D}_h^T\mathbf{D}_h$ is similar as defined above and the matrix $\mathbf{D}_v^T\mathbf{D}_v$ is:

$$\mathbf{D}_v^T\mathbf{D}_v = \begin{pmatrix} 1 & -2 & 1 & & & & \\ -2 & 5 & -4 & 1 & & & \\ 1 & -4 & 6 & -4 & 1 & & \\ & 1 & -4 & 6 & -4 & 1 & \\ & & 1 & -4 & 6 & -4 & 1 \\ & & & 1 & -4 & 5 & -2 \\ & & & & 1 & -2 & 1 \end{pmatrix}.$$

Thus, we have:

$$\mathbf{I}_4 \otimes (\mathbf{D}_v^T\mathbf{D}_v) = \begin{array}{c} \\ \cdot \\ \cdot \\ \cdot \\ \gamma_{11} \\ \gamma_{12} \\ \gamma_{13} \\ \gamma_{14} \\ \vdots \\ \gamma_{41} \\ \gamma_{42} \\ \gamma_{43} \\ \gamma_{44} \\ \cdot \\ \cdot \\ \cdot \end{array} \begin{array}{c} \begin{array}{ccccccccccccccc} \cdot & \cdot & \cdot & \gamma_{11} & \gamma_{12} & \gamma_{13} & \gamma_{14} & \cdots & \gamma_{41} & \gamma_{42} & \gamma_{43} & \gamma_{44} & \cdot & \cdot & \cdot \end{array} \\ \begin{pmatrix} 1 & -2 & 1 & & & & & & & & & & & & \\ -2 & 5 & -4 & 1 & & & & & & & & & & & \\ 1 & -4 & 6 & -4 & 1 & & & & & & & & & & \\ & 1 & -4 & 6 & -4 & 1 & & & & & & & & & \\ & & 1 & -4 & 6 & -4 & 1 & & & & & & & & \\ & & & 1 & -4 & 5 & -2 & & & & & & & & \\ & & & & 1 & -2 & 1 & & & & & & & & \\ & & & & & & & \ddots & & & & & & & \\ & & & & & & & & 1 & -2 & 1 & & & & \\ & & & & & & & & -2 & 5 & -4 & 1 & & & \\ & & & & & & & & 1 & -4 & 6 & -4 & 1 & & \\ & & & & & & & & & 1 & -4 & 6 & -4 & 1 & \\ & & & & & & & & & & 1 & -4 & 6 & -4 & 1 \\ & & & & & & & & & & & 1 & -4 & 5 & -2 \\ & & & & & & & & & & & & 1 & -2 & 1 \end{pmatrix} \end{array}$$

and

$$
(\mathbf{D}_h^T\mathbf{D}_h)\otimes\mathbf{I}_7 = \begin{array}{c} \\ \cdot \\ \cdot \\ \cdot \\ \gamma_{11} \\ \gamma_{12} \\ \gamma_{13} \\ \gamma_{14} \\ \cdot \\ \cdot \\ \gamma_{21} \\ \gamma_{22} \\ \gamma_{23} \\ \gamma_{24} \\ \cdot \\ \vdots \end{array}
\begin{array}{c}
\begin{array}{ccccccccccccccc}
\cdot & \cdot & \cdot & \gamma_{11} & \gamma_{12} & \gamma_{13} & \gamma_{14} & \cdot & \cdot & \gamma_{21} & \gamma_{22} & \gamma_{23} & \gamma_{24} & \cdot & \cdots
\end{array} \\
\left(\begin{array}{ccccccccccccccc}
1 & & & & & & & -2 & & & & & & & \ddots \\
 & 1 & & & & & & & -2 & & & & & & \\
 & & 1 & & & & & & & -2 & & & & & \\
 & & & 1 & & & & & & & -2 & & & & \\
 & & & & 1 & & & & & & & -2 & & & \\
 & & & & & 1 & & & & & & & -2 & & \\
 & & & & & & 1 & & & & & & & -2 & \\
-2 & & & & & & & 5 & & & & & & & \ddots \\
 & -2 & & & & & & & 5 & & & & & & \\
 & & -2 & & & & & & & 5 & & & & & \\
 & & & -2 & & & & & & & 5 & & & & \\
 & & & & -2 & & & & & & & 5 & & & \\
 & & & & & -2 & & & & & & & 5 & & \\
 & & & & & & -2 & & & & & & & 5 & \\
\ddots & & & & & & & \ddots & & & & & & & \ddots
\end{array}\right)
\end{array}.
$$

In Section 2.3 all matrices and vectors are similarly defined as in Section 2.1. In the penalty matrix $\mathbf{P}$ in (12), the matrix $\mathbf{D}_h^T\mathbf{D}_h$ is similar as defined above. The example, in the case where $m = 4$, of the penalty matrix $P_d$ is presented in the manuscript.

### Software Code

R version 4.1.2 is used to fit the proposed models. We make use of sparse matrix implementations using the R-package Matrix.

**Software Code for the Construction of the Penalty Matrices**

R-code for the penalty $\mathbf{P}$ in (5):

```
Dh <- diff(Diagonal(m), diff=2)
Dv <- diff(Diagonal(m), diff=2)
Ph <- kronecker(Diagonal(m), t(Dh)%*%Dh)
Pv <- kronecker(t(Dv)%*%Dv, Diagonal(m))
P <- lambda1*Ph + lambda2*Pv
```

R-code for the penalty $\mathbf{P}$ in (11):

```
mm <- 2*m-1
Dv <- diff(Diagonal(mm), diff=2)
Dh <- diff(Diagonal(m), diff=2)
Pv <- kronecker(Diagonal(m), t(Dv)%*%Dv)
Ph <- kronecker(t(Dh)%*%Dh, Diagonal(mm))
P <- lambda1*Pv + lambda2*Ph
```

For the construction of the penalty $\mathbf{P}$ in (12) an algorithm is needed to construct the penalty matrix $\mathbf{P}_d$:

```
Dh = diff(Diagonal(m), diff=2)
Ph = kronecker(Diagonal(m) , t(Dh)%*%Dh)
Pd = Matrix(0,nrow=m*m,ncol=m*m)
### Algorithm to construct Pd
for (i in 1:m){
  for (j in 1:m){
    dim.input = m-abs(j-i)
    if (dim.input > 2){
      D <- diff(Diagonal(dim.input), diff=2)
      DD <- t(D) %*% D
    }
    if (dim.input == 2){
      D <- diff(Diagonal(2), diff=1)
      DD <- t(D) %*% D
    }
    if (dim.input == 1){
      DD <- matrix(0,nrow=1,ncol=1)
    }
    if ( i <= j ){
      index1 <- m*(i-1) + j
      index2 <- seq( ((j-i)+1) , (m*(m-(j-i))) , (m+1) )
      Pd[index1,index2] <- DD[i,]
    }
    if ( i > j ){
      index1 <- m*(i-1) + j
      index2 <- seq( (1+m*(i-j)) , (m*m-(i-j)) , (m+1) )
      Pd[index1,index2] <- DD[j,]
    }
  }
} # end of algorithm
P <- lambda1*Ph + lambda2*Pd
```

For the construction of the penalty **P** in (11) with a kink on the main diagonal, we use:

```
tilde.I = Diagonal(mm)
tilde.I[mm/2+0.5,mm/2+0.5] = 0
Dv.star <- diff(tilde.I, diff=2)
Dv.star[(m-2),(m-1)]= -1
Dv.star[(m-1),(m+1)]= -1
Dv.star[m,(m+1)]= -1
Pv <- kronecker(Diagonal(x=c(rep(1,max.kink.age),rep(0,m-max.kink.age))),
      t(Dv.star)%*%Dv.star)
      + kronecker(Diagonal(x=c(rep(0,mka),rep(1,n-mka))),
      t(Dxx1.nonadj)%*%Dv)
```

For the construction of the penalty **P** in (14) with a kink on the main diagonal, the following code is used to construct the penalty in the horizontal dimension:

```
Ph = Matrix(0,nrow=m*m, ncol=n*n)
matrix.kink.adjustment = t(Matrix(c(1,-1,0,0,0, 0,1,0,-1,0, 0,0,0,-1,1),5,3))
for (i in 1:n){
  Dh  = diff(Diagonal(n), diff=2)
  if (i==1 & i<max.kink.age){
    Dh[1,1:3] = matrix.kink.adjustment[-c(1:2),-c(1:2)]
  }
  if (i==2 & i<max.kink.age){
    Dh[1:2,1:4] = matrix.kink.adjustment[-c(1),-c(1)]
  }
  if (i==(n-1) & i<max.kink.age){
    Dh[(n-3):(n-2) , (n-3):n] = matrix.kink.adjustment[-c(3) , -c(5)]
  }
  if (i==n & i<max.kink.age){
    Dh[(n-2) , (n-2):n] = matrix.kink.adjustment[-c(2:3) , -c(4:5)]
  }
  if (i %in% (3:(n-2)) & i<max.kink.age){
    Dh[((i-2):(i)) , ((i-2):(i+2))] = matrix.kink.adjustment
  }
  Ph[((n*(i-1)+1):(n*i)) , ((n*(i-1)+1):(n*i))] = t(Dh)%*%Dh
}
```

#### Software Code for the Reordening of the Contact Matrix

To construct the restructured matrices $\breve{\mathbf{Y}}$, $\breve{\mathbf{E}}$ and $\breve{\boldsymbol{\Gamma}}$ the following R-code can be used:

```
mm <- 2*m-1
Pos <- cbind(c(row(Y)-col(Y)+m), c(col(Y)))
Y.tilde <- matrix(NA,mm,m)
Y.tilde[Pos] <- c(t(Y))
```

The construction of the matrix $\breve{\mathbf{W}}$ can be done using:

```
W.tilde <- matrix(0, mm, m)
W.tilde[Pos] <- 1
```

#### Software Code for the Constrained Penalized Iterative Reweighted Least Squares (C-PIRLS) Estimation

The R-code of the C-PIRLS scheme in the case of a Poisson distribution using the methodology of Section 2.1 is given first. We remark once more that parameters estimation using penalized iterative reweighted least squares without using the symmetry constraint is done first and use the obtained estimated parameters as starting values in the C-PIRLS fitting.

```
### Starting values
eta <- Matrix(0, m*m, 1)
eta <- log((y+1)/(e+1))

### PIRLS (without symmetry constraint)
for (it in 1:max.iter){
  mu.vector = e * exp(eta)
  mu <- Matrix(mu.vector, m*m, 1)
  W <- Diagonal(x=as.vector(mu))
  pseudo <- eta + 1/mu*(y-mu)
  r <- W %*% pseudo
  WP <- W + P
  etanew <- solve(WP, r)
  deta <- max(abs(etanew - eta))
  eta <- etanew
  if(deta<=10^-4 & it>=3) break
}

### Construct the constraint matrix L and vector nu
posY <- col(Y) - row(Y)
NU0 <- matrix(log(p), m, m)
NU1 <- t(NU0)
NU01 <- NU0-NU1
nu <- c(NU01[which(posY<0)])
nc <- length(nu) # number of constraints
L <- Matrix(0, nc, m*m)
pos0 <- rep(1:m, m)
pos1 <- rep(1:m, each=m)
  # placing 1
cc1a <- outer(seq(0, by=m, length=m), seq(1, m), "+")
cc1 <- cc1a[row(cc1a) < col(cc1a)]
rr1 <- 1:nc
L[cbind(rr1,sort(cc1))] <- 1
  # placing -1
cc2a <- outer(seq(0, by=m, length=m), seq(1, m), "+")
cc2 <- cc2a[row(cc2a) > col(cc2a)]
rr2 <- 1:nc
L[cbind(rr2,cc2)] <- -1

### C-PIRLS
for (it in 1:max.iter){
  mu.vector <- e * exp(eta)
  mu <- Matrix(mu.vector, m*m, 1)
  W <- Diagonal(x=as.vector(mu))
  pseudo <- eta + 1/mu*(y-mu)
  r <- W %*% pseudo
  WP <- W + P
  LHS <- Matrix(0, m*m+nc, m*m+nc)
  LHS[1:(m*m),1:(m*m)] <- WP
  LHS[1:(m*m),1:nc+(m*m)] <- t(L)
  LHS[1:nc+(m*m), 1:(m*m)] <- L
  RHS <- Matrix(0, (m*m)+nc, 1)
  RHS[1:(m*m)] <- r
  RHS[1:nc+(m*m)] <- nu
  eta.ome <- solve(LHS, RHS)
  etanew <- eta.ome[1:(m*m)]
  deta <- max(abs(eta - etanew))
  eta <- etanew
  if(deta<=10^-4 & it>=3) break
}

### -2*log-likelihood
mu.ll <- exp(eta) * e
ll <- -2*sum( y*log(mu.ll) - mu.ll - lfactorial(y) )
### Estimated ETA and GAMMA matrices
ETA.S <- t(matrix(eta, m, m))
GAMMA.S <- exp(ETA.S)
### Effective degrees of freedom & AIC
H = solve(WP,W)
edf = sum(diag(H))
aic = ll + 2*(edf)
```

In case the negative binomial distribution is of interest, the following R-code is used for the PIRLS without symmetry constraint:

```
### PIRLS (without symmetry constraint)
phi=0.00
for (it1 in 1:max.iter.phi){
  for (it in 1:max.iter){
    mu.vector = e * exp(eta)
    mu <- Matrix(mu.vector, m*m, 1)
    W <- Diagonal( x=as.vector(mu)/(1+phi) )
    pseudo <- eta + 1/mu*(y-mu)
    r <- W %*% pseudo
    WP <- W + P
    etanew <- solve(WP, r)
    deta <- max(abs(etanew - eta))
    eta <- etanew
    if(deta<=10^-4 & it>=3) break
  }
  # Estimate dispersion parameter phi
  H = solve(WP,W)
  edf = sum(diag(H))
  phi.old = phi
  mu.fit = exp(eta) * e
  phi.new = sum(1/mu.fit * (y-mu.fit)^2)/(m*m - edf) - 1
  diff.phi = abs(phi.new - phi.old)
  phi = phi.new
  if (diff.phi<=10^-4) break
}
```

For the part of the C-PIRLS fitting the following R-code is used:

```
### C-PIRLS
phi = phi.input
for (it1 in 1:max.iter.phi){
  for (it in 1:max.iter){
    mu.vector <- e * exp(eta)
    mu <- Matrix(mu.vector, m*m, 1)
    W <- Diagonal( x=as.vector(mu)/(1+phi) )
    pseudo <- eta + 1/mu*(y-mu)
    r <- W %*% pseudo
    WP <- W + P
    LHS <- Matrix(0, m*m+nc, m*m+nc)
    LHS[1:(m*m),1:(m*m)] <- WP
    LHS[1:(m*m),1:nc+m*m] <- t(L)
    LHS[1:nc+(m*m), 1:(m*m)] <- L
    RHS <- Matrix(0, (m*m)+nc, 1)
    RHS[1:(m*m)] <- r
    RHS[1:nc+(m*m)] <- nu
    eta.ome <- solve(LHS, RHS)
    etanew <- eta.ome[1:(m*m)]
    deta <- max(abs(eta - etanew))
    eta <- etanew
    if(deta<=10^-4 & it>=3) break
  }
  # Estimate dispersion parameter phi
  H = solve(WP,W)
  edf = sum(diag(H))
  phi.old = phi
  mu.fit = exp(eta) * e
  phi.new = sum(1/mu.fit * (y-mu.fit)^2)/(m*m - edf) - 1
  diff.phi = abs(phi.new - phi.old)
  phi = phi.new
  if (diff.phi<=10^-4) break
}
  # final estimation of eta
for (it in 1:max.iter){
  mu.vector <- e * exp(eta)
  mu <- Matrix(mu.vector, m*m, 1)
```

```
    W <- Diagonal( x=as.vector(mu)/(1+phi) )
    pseudo <- eta + 1/mu*(y-mu)
    r <- W %*% pseudo
    WP <- W + P
    LHS <- Matrix(0, m*m+nc, m*m+nc)
    LHS[1:(m*m),1:(m*m)] <- WP
    LHS[1:(m*m),1:nc+m*m] <- t(L)
    LHS[1:nc+(m*m), 1:(m*m)] <- L
    RHS <- Matrix(0, (m*m)+nc, 1)
    RHS[1:(m*m)] <- r
    RHS[1:nc+(m*m)] <- nu
    eta.ome <- solve(LHS, RHS)
    etanew <- eta.ome[1:(m*m)]
    deta <- max(abs(eta - etanew))
    eta <- etanew
    if(deta<=10^-4 & it>=3) break
  }

  ### -2*log-likelihood
  mu.ll = exp(eta) * e
  ll = -2*sum( lfactorial(y + mu.ll/phi - 1) - lfactorial(y+1-1) -
    lfactorial(mu.ll/phi - 1) + mu.ll/phi*log((mu.ll/phi)/(mu.ll/phi+mu.ll)) +
    y*log((mu.ll)/(mu.ll/phi+mu.ll)) )
  ### Estimated ETA and GAMMA matrices
  ETA.S <- t(matrix(eta, m, m))
  GAMMA.S <- exp(ETA.S)
  ### Effective degrees of freedom & AIC
  H = solve(WP,W)
  edf = sum(diag(H))
  aic = ll + 2*(edf+1)
```

The software code to fit the models described in Section 2.3 is similar as the software code used for the models in Section 2.1. To implement the methodology described in Section 2.2, some modifications are needed to the R-code. The R-code for the Poisson distribution is presented here:

```
  ### Starting values
  pos <- which(W.tilde==1)
  mm <- 2*m-1
  mmm <- mm*m
  eta <- Matrix(0, mmm, 1)
  eta[pos] <- log((tilde.y+1)/(tilde.e+1))[pos]

  ### PIRLS (without symmetry constraint)
  for (it in 1:max.iter){
    mu <- Matrix(0, mmm, 1)
    mu[pos] <- tilde.e[pos] * exp(eta)[pos] * w.tilde[pos]
    W <- Matrix(0, mmm,mmm)
    W[cbind(pos, pos)] <- mu@x
    pseudo <- Matrix(0, mmm, 1)
    pseudo[pos] <- eta[pos] + 1/mu@x*(tilde.y[pos] - mu@x)
    r <- W %*% pseudo
    WP <- W + P
    etanew <- solve(WP, r)
    deta <- max(abs(etanew - eta))
    eta <- etanew
    if(deta<=10^-4 & it>=3) break
  }

      ### Construct the constraint matrix L and vector nu
  NU0 <- matrix(log(p), m, m)
  NU1 <- t(NU0)
  NU01 <- NU0-NU1
  nu <- c(NU01[which(posY<0)])
  nc <- length(nu) ## number of constraints

  L <- Matrix(0, nc, mmm)
  pos0 <- rep(1:mm, m)
  pos1 <- rep(1:m, each=mm)
    # placing 1
```

```
cc1a <- outer(seq(m+1, mm), seq(0, by=mm, length=m-1), "+")
cc1 <- cc1a[row(cc1a)+col(cc1a)<=m]
rr1 <- 1:nc
L[cbind(rr1,cc1)] <- 1
  # placing -1
cc2a <- outer(seq(0, by=mm-1, length=m-1), seq(mm+m-1, length=m-1,by=mm), "+")
cc2 <- cc2a[row(cc2a)+col(cc2a)<=m]
rr2 <- 1:nc
L[cbind(rr2,cc2)] <- -1

### C-PIRLS
for (it in 1:max.iter){
  mu <- Matrix(0, mmm, 1)
  mu[pos] <- tilde.e[pos] * exp(eta)[pos] * w.tilde[pos]
  W <- Matrix(0, mmm,mmm)
  W[cbind(pos, pos)] <- mu@x
  pseudo <- Matrix(0, mmm, 1)
  pseudo[pos] <- eta[pos] + 1/mu@x*(tilde.y[pos] - mu@x)
  r <- W %*% pseudo
  WP <- W + P
  LHS <- Matrix(0, mmm+nc, mmm+nc)
  LHS[1:mmm,1:mmm] <- WP
  LHS[1:mmm,1:nc+mmm] <- t(L)
  LHS[1:nc+mmm, 1:mmm] <- L
  RHS <- Matrix(0, mmm+nc, 1)
  RHS[1:mmm] <- r
  RHS[1:nc+mmm] <- nu
  eta.ome <- solve(LHS, RHS)
  etanew <- eta.ome[1:mmn]
  deta <- max(abs(eta - etanew))
  eta <- etanew
  if(deta<=10^-4 & it>=3) break
}

### -2*log-likelihood
y.ll = tilde.y[pos]
mu.ll = exp(eta[pos]) * tilde.e[pos]
ll = -2*sum( y.ll*log(mu.ll) - mu.ll - lfactorial(y.ll) )
### Estimateed ETA and GAMMA matrices
eta.hat.vector <- rep(NA,mmm)
eta.hat.vector[pos] <- eta[pos]
eta.hat.matrix <- matrix(eta.hat.vector, mm, m)
ETA.S <- matrix(NA, m, m)
for(i in 1:m){
  ETA.S[,i] <- eta.hat.matrix[!is.na(eta.hat.matrix[,i]), i]
}
ETA.S <- t(ETA.S)
GAMMA.S <- exp(ETA.S)
### Effective degrees of freedom
H = solve(WP,W)
edf = sum(diag(H))
aic = ll + 2*(edf)
```

This code can be adjusted as above for the negative Binomial distribution.
